# SPAFESTWDILK, a plant-derived dodecapeptide from *Zingiber officinale*, as a predicted inhibitor of the MDM2–p53 interaction: computational discovery and multi-method evaluation

**DOI:** 10.64898/2026.06.06.730565

**Authors:** Massimiliano Romiti, Carla Sandri, Giulia Paiola, Minoo Ashtiani

## Abstract

The MDM2-p53 protein-protein interaction is a validated oncology target, yet no food-derived linear peptide has been documented to engage the canonical three-anchor MDM2-p53 interface. We developed a multi-stage computational pipeline (PepVeg) to screen 22 plant and fungal proteomes (337,646 proteins) for MDM2-binding peptides, applying sequential in silico hydrolysis, physicochemical filtering, ESM-2 embedding-based dimensionality reduction, and pharmacophore-driven selection. Twenty-six candidates were evaluated by AlphaFold 3 (AF3) co-folding against MDM2(25-109), yielding 15 binders (iPTM >= 0.75; 58% of evaluated). A 36-peptide benchmark with 29 hard negatives confirmed AF3 discriminative power (Cohen’s d = 3.41; 95% CI: 1.94-4.88; Hedges’ g = 3.32; zero overlap).

The lead candidate, SPAFESTWDILK -- a tryptic fragment of Zingiber officinale histone deacetylase (UniProt A0A8J5FLH2) -- was evaluated by eight computational assessments: AF3 Server (iPTM 0.83, SD 0.01), Protenix (iPTM 0.923), Chai-1 (iPTM 0.891), EvoEF2 (−55.57 EEU), two GROMACS simulations (no dissociation across two force fields), and two MM-PBSA calculations (−75.30 (SD 4.92) and −55.07 (SD 2.86) kcal/mol). The W8A point mutant produced an iPTM drop of 0.201, closely paralleling the p53 W23A drop of 0.193; we predict W8A substitution will abolish binding. SPAFESTWDILK ranked only #890/2,000 by ESM-2 similarity and was recovered solely through pharmacophore matching, demonstrating that no single pipeline stage alone is sufficient. To our knowledge, this is the first food-database-derived linear peptide with multi-convergent computational evidence supporting engagement of the canonical three-anchor MDM2-p53 interface. Experimental validation by SPR/ITC is warranted.

## 1. INTRODUCTION

The tumor suppressor p53 is inactivated in over half of human cancers, frequently through overexpression of its negative regulator MDM2 [1]. The co-crystal structure of the MDM2 N-terminal domain with the p53 transactivation domain revealed a deep hydrophobic cleft into which three p53 residues — Phe19, Trp23, and Leu26 — insert as sub-pocket anchors [1]. This F-W-L triad is the structural fingerprint of every known MDM2-binding peptide, including PMI (Kd = 3.3 nM [2]), 12/1 (Kd ∼200 nM [3]), the native p53(17-29) (Kd ∼580 nM [20]), and p53(15-29) (Kd = 140 nM [2]). The two p53 fragments differ in length: p53(17-29) (13 residues) and p53(15-29) (15 residues, including additional N-terminal contacts); the shorter fragment was used in the AF3 calibration panel, the longer in the Protenix discrimination panel. Despite extensive pharmacology, the principal MDM2 inhibitor series is synthetic: Nutlin derivatives, phage-display peptides, and stapled helices [4,5,23,24]. Among natural products, chlorofusin — a cyclic peptide from the fungus *Microdochium caespitosum* — has been reported to inhibit the MDM2–p53 interaction (IC50 ∼4.7 μM) [21], but does not engage the three-anchor p53-mimicking interface. To our knowledge, no food-derived linear peptide has been documented to target the canonical Phe19-Trp23-Leu26 MDM2–p53 binding interface.

Despite extensive pharmacological effort, the principal MDM2 inhibitor series remains synthetic. Nutlin-3a demonstrated proof-of-concept for pharmacological p53 reactivation *in vivo* [23], and nine small-molecule candidates have entered clinical evaluation since, none of which has yet received regulatory approval, partly due to on-target haematological toxicity and resistance acquired through TP53 mutation [24,37]. Stapled alpha-helical peptides represent a complementary approach: by introducing hydrocarbon crosslinks that stabilise helical conformation, dual MDM2/MDMX inhibitors achieve sub-nanomolar potency while retaining cell permeability [4,5]. Among natural products, chlorofusin — a cyclic depsipeptide from *Microdochium caespitosum* — inhibits MDM2–p53 with IC50 ∼4.7 µM [21], but does not engage the canonical three-anchor p53-mimicking interface. Alternative scaffolds with distinct physicochemical profiles, including food-derived peptides, are therefore of continued interest [31].

Recent advances in protein language models (pLMs) provide a practical solution to proteome-scale screening. ESM-2, trained by masked language modelling on 250 million sequences [9], generates embeddings that capture evolutionary context at amino acid resolution. ESM-2 embeddings have been applied as pre-filters for antihypertensive [29] and antimicrobial peptide discovery [30], and PepMLM — a derivative model fine-tuned on protein–peptide interaction data — has generated potent MDM2-binding sequences de novo without structural input [31]. Complementing embedding-based screening, AlphaFold 3 (AF3) predicts protein–peptide complex geometries with high accuracy for helical ligands in defined grooves, as established by large-scale benchmarking [17,32,33]; the interface predicted transfer metric (iPTM) provides a per-complex confidence score correlated with structural correctness at the interface.

Plant proteomes are an established source of bioactive peptides [6]. *Zingiber officinale* (ginger) yields tryptic peptides with antioxidant and ACE-inhibitory activities [7,8,40], but none has been screened against the MDM2–p53 interaction. Screening a large proteome corpus (≥ 10⁵ proteins) for MDM2 binders by direct structural prediction is computationally prohibitive. We therefore developed a cascaded pipeline combining *in silico* hydrolysis, physicochemical filtering, ESM-2 embedding similarity, pharmacophore-driven selection, and AF3 co-folding, followed by independent multi-method evaluation.

The objectives of this study are: (1) to describe the pipeline and characterize its performance on MDM2; (2) to identify and computationally evaluate SPAFESTWDILK as a candidate MDM2 binder through eight computational assessments; and (3) to present the W8A point mutation result as a directly testable prediction.

## 2. MATERIALS AND METHODS

### 2.1 Target pre-screening and construct optimization

The pipeline includes a target tractability assessment prior to peptide screening. AI co-folding tools achieve high accuracy for helical peptides in defined grooves [32,33] but perform poorly on flat-interface or multi-subunit complexes [44,45]. Five criteria were applied to evaluate target suitability: (i) minimal binding domain: ≤ 200 residues (avoids disordered regions confounding AF3 confidence); (ii) deep binding pocket: BSA ≥ 400 Å², groove geometry (not flat interface); (iii) validated binders with Kd: ≥ 3 binders spanning > 100-fold affinity range (enables calibration); (iv) known pharmacophore: linear motif enabling regex-based selection from peptide pools; and (v) defined mechanism of action: inhibition or activation (determines screening logic). MDM2 passes all five criteria: 85-residue monomer (UniProt Q00987, residues 25–109), deep hydrophobic cleft (∼700 Å² BSA [1,22]), 5 validated binders spanning ∼176-fold (175.8×) (PMI 3.3 nM to p53(17-29) ∼580 nM), the F19-W23-L26 pharmacophore triad, and competitive inhibition of the MDM2–p53 interaction. Cross-target comparison with EphA2 (moderate tractability: 174 aa, one binder at ∼1 μM, weak pharmacophore) and IL-11 (low tractability: flat interface, no binders with Kd, no pharmacophore) confirmed MDM2 as the highest-priority target. Empirical validation: AF3 iPTM gap between binders and non-binders was 0.248 on MDM2 vs 0.103 on IL-11 controls [a,d].

The p53-binding cleft is formed by 14 key residues organized into three sub-pockets: the Phe19 pocket (Ile61, Met62, Tyr67, Val75, Gln72), the Trp23 pocket (Leu54, Leu57, Gly58, Phe86, Phe91, Val93, Ile99, Ile101 — deepest sub-pocket, indole N-H forms H-bond to Leu54 backbone carbonyl at ∼2.9 Å [22]), and the Leu26 pocket (Leu54, Val93, His96, Ile99, Tyr100). The pharmacophore F-x(3)-W places F and W at alpha-helix positions i and i+4, projecting both side chains into adjacent sub-pockets. This structural arrangement determines which peptides from the screening pool will achieve correct three-pocket occupancy in AF3 co-folding.

The 85-residue minimal binding domain was used throughout, based on a direct comparison showing that removal of 24 disordered N-terminal residues improved AF3 iPTM from 0.57 to 0.85 for SPAFESTWDILK while preserving inter-peptide discrimination (0.11 iPTM gap between candidates) [d]. This improvement reflects better tool calibration, not independent validation of binding.

### 2.2 AF3 calibration panel

AF3 was calibrated on a 3+3 panel of three validated binders (PMI, 12/1, p53(17-29)) and three non-binders (W23A, poly-Ala, scrambled p53) before candidate screening. Complete separation was observed (minimum gap +0.248 iPTM), establishing provisional working thresholds: iPTM ≥ 0.75 (binder), < 0.65 (non-binder). We note that n = 6 is insufficient for robust threshold derivation; these are working decision boundaries supplemented by the downstream expanded benchmark (Section 2.6).

### 2.3 Proteome screening pipeline

#### Step 1: Hydrolysis

Twenty-two proteomes (337,646 proteins) from edible, medicinal, and food-waste sources were subjected to *in silico* enzymatic hydrolysis with trypsin, bromelain, and pepsin (pH > 2), retaining fragments of 7–12 residues (8,645,509 unique peptides) [a].

#### Step 2: Physicochemical filtering

A heuristic pre-filter (Boman index ≥ 1.0 [18], GRAVY −2.0 to +0.5, net charge −2 to +4, instability index < 40, MW < 1,500 Da, hydrophobicity 30–60%) reduced the pool to 1,642,848 peptides (−81.0%) [a]. Filters were applied sequentially in the order listed; each removal count reflects peptides eliminated from the pool remaining after all prior filters. This step constitutes chemical space reduction, not biologically validated selection; the cumulative filters introduce a bias toward canonical PPI-targeting chemotypes. Full filter specifications are available in Supplementary Table S2.

Two implementation notes on the pre-filter are relevant to the lead candidate. First, the Boman index was computed using a modified Radzicka–Wolfenden transfer energy scale (see pipeline source code); SPAFESTWDILK scores 1.24 on this scale [a], passing the ≥ 1.0 threshold. Second, the instability index was computed using a simplified DIWV dipeptide weight matrix covering a subset of the 400 possible dipeptides (unmatched dipeptides default to a weight of 1.0). Under this implementation, SPAFESTWDILK scores −3.36 and passes the < 40 threshold [a]. The standard ProtParam implementation (complete 400-entry DIWV matrix) yields 74.33 [z], which would not pass the filter. This discrepancy reflects the incomplete DIWV matrix in the pipeline and is disclosed here for transparency. We note that the Guruprasad instability index was calibrated on full-length proteins and is not validated for short peptides (12 residues); neither the pipeline value nor the ProtParam value should be interpreted as a reliable stability prediction for a dodecapeptide.

#### Step 3: ESM-2 dimensionality reduction

Peptides were embedded using ESM-2 (esm2_t33_650M_UR50D [9,29,30]). Cosine similarity to p53(17-29) was computed, and the top 2,000 peptides were retained [b]. ESM-2 cosine similarity shows near-zero correlation with binding affinity (Pearson r = −0.049 on 97 IL-11 peptides [a]) and serves as dimensionality reduction, not affinity ranking. The top-2,000 cutoff is a practical heuristic; its sensitivity has not been formally evaluated.

#### Step 4: Pharmacophore selection

For MDM2, the regex F.{1,3}W was applied, retaining 81 peptides (4.0% of 2,000) [c]. A manual scoring scheme (0–4.5) prioritized AF3 submission order but is not predictive of binding (Section 3.3).

#### Step 5: AF3 screening

Twenty-six candidates were evaluated on AF3 Server with the 85-residue MDM2 construct, generating 5 models each. Candidates were prioritized for AF3 evaluation based on: (1) length 10–14 residues (matching PMI/p53 reference range), (2) species diversity (max 3 per species), (3) pharmacophore score ≥ 2.5/4.5. All 26 evaluated candidates met these criteria prospectively. The 58% hit rate is applicable to this pre-filtered subset, not to all 81 pharmacophore-matched candidates.

#### Step 6: GROMACS triage

AF3 hits underwent 5 ns GROMACS triage (CHARMM36m [38]; pipeline code available at https://github.com/Kingelanci/PepVeg-Pipeline) as a coarse instability filter; candidates with RMSD < 0.25 nm were promoted to 100 ns production runs [n].

### 2.4 Independent multi-method evaluation

SPAFESTWDILK (AF3 Server iPTM best 0.85, mean 0.83 ± 0.01) was independently evaluated on the Neurosnap platform using: (1) Protenix (AF3 reproduction, 5 models); (2) Chai-1 (5 seeds); (3) EvoEF2 ComputeBinding [10]; (4) GROMACS 50 ns (AMBER99SB-ILDN [11], TIP3P, 0.15 M NaCl, 310 K); (5) MM-PBSA (gmx_MMPBSA [12]) on the 50 ns trajectory. A second MM-PBSA was computed on a CHARMM27 50 ns trajectory. Cross-method agreement reflects robustness of a single binding hypothesis under diverse approximations, not independent thermodynamic validation, as the structural inputs share a common origin. Full Neurosnap job parameters and share URLs for all platform analyses are listed in Supplementary Table S1.

### 2.5 Seven-peptide discrimination panel

A panel was run on Protenix with identical parameters: PMI, p53(15-29), SPAFESTWDILK (binders), and poly-Ala, p53 W23A, SPAFESTWDILK W8A (SPAFESTADILK), scrambled (non-binders). Effect sizes (Cohen’s d, Hedges’ g) and 95% confidence intervals were computed using df-weighted pooled SD. Full derivation of effect size statistics for both panels, including individual data, pooling formula, confidence intervals, and Hedges’ g correction, is provided in Supplementary Table S7. With n = 3 vs n = 4, Cohen’s d is provided as a descriptive summary and not as a basis for formal statistical inference.

### 2.6 Expanded benchmark (29 hard negatives)

To address the small-N limitation of the 3+3 panel, 29 hard negatives were evaluated across four categories: (1) W-position variants (n = 8): SPAFESTWDILK with Trp relocated to non-native positions, composition-matched, lengths 11–12; (2) scrambled no-W (n = 5): W→A substitution with full scrambling, length 12; (3) hydrophobic no-W (n = 8): W replaced by other hydrophobic residues, lengths 11–12; (4) proteome-random (n = 8): tryptic peptides from housekeeping proteins, filter-matched only, lengths 10–14. All tested on AF3 with 5 seeds each against MDM2(25–109). Length was not strictly controlled across categories. Combined with 3 binders and 4 original controls: 36-peptide panel. Per-seed data are available in Supplementary Table S3.

### 2.7 Precursor identification

The parent protein was identified by BLAST (exact substring match: A0A8J5FLH2, *Zingiber officinale* histone deacetylase, PE = 3). Ginger rhizome transcriptomics confirms HDAC-family transcript expression [25,26]. Simulated GI digestion was performed with ExPASy PeptideCutter [13].

## 3. RESULTS

### 3.1 Pipeline funnel (Fig. S1)

The six-stage PEPVEG pipeline processed 337,646 proteins from 22 plant and fungal proteomes, generating 8,645,509 unique tryptic, bromelain, and pepsin fragments (7–12 residues) (Table 1). Sequential physicochemical filtering (Boman index, GRAVY, net charge, instability index, MW, hydrophobic fraction) reduced the pool by 81.0% to 1,642,848 peptides. ESM-2 cosine similarity ranking against p53(17-29) retained the top 2,000 sequences, followed by pharmacophore-driven selection (regex F.{1,3}W), yielding 81 candidates (4.0% of 2,000). Twenty-six candidates meeting length, species diversity, and pharmacophore score criteria were submitted to AF3 co-folding against MDM2(25–109); 15 (58%) achieved iPTM ≥ 0.75. GROMACS 5 ns triage then reduced these to 3 candidates for full multi-method evaluation. Notably, SPAFESTWDILK ranked only #890/2,000 by ESM-2 cosine similarity and was recovered exclusively through the pharmacophore filter (Stage 4), demonstrating that no single pipeline stage alone is sufficient (Fig. S1)

**Table 1.** Pipeline funnel summary. Data references in brackets correspond to deposited files at Zenodo DOI: 10.5281/zenodo.19859570.

| Step | Filter | Output | Reduction | Ref |
| --- | --- | --- | --- | --- |
| 1 | 22 proteomes (337,646 proteins), in silico hydrolysis | 8,645,509 peptides | — | [a] |
| 2 | Physicochemical pre-filter | 1,642,848 | −81.0% | [a] |
| 3 | ESM-2 top 2,000 (MDM2) | 2,000 | −99.9% | [b] |
| 4 | Pharmacophore F.<sub>{1,3}</sub>W | 81 | −96% | [c] |
| 5 | AF3 screening (26 evaluated) | 15 binders (iPTM $\geq 0.75$) | 58% of evaluated | [d] |
| 6 | EvoEF2 + GROMACS triage | 3 candidates | final | [h,n] |

### 3.2 AF3 calibration

The 3+3 panel yielded complete separation between binders and non-binders on MDM2 (Table 2). AF3 iPTM ranked binders consistently with experimental affinity: PMI (0.890) > 12/1 (0.870) > p53(17-29) (0.830) [d]. The W23A mutant scored iPTM avg 0.532 (best 0.640, SD 0.096) [d], a drop of 0.298 from wild-type p53 avg (0.830). The high W23A variance (SD 0.096 vs SD 0.007 for binders) is itself informative: the loss of the Trp23 anchor destabilizes the predicted pose. In a parallel assessment, PRODIGY failed to separate binders from non-binders on MDM2 [31] (scrambled peptide scored above 4/6 binders), consistent with its calibration on globular complexes rather than short peptide-groove systems.

**Table 2.** AF3 calibration panel (MDM2). All values verified from raw AF3 Server JSON files [d] (5 models per peptide, MDM2 85-residue construct). SPAFESTWDILK grand mean 0.828 ± 0.011 (10 seeds).

| Peptide | Class | Kd | iPTM avg<br>(5 seeds) | iPTM<br>best | SD | Ref |
| --- | --- | --- | --- | --- | --- | --- |
| PMI | binder | 3.3 nM | 0.890 | 0.900 | 0.007 | [d] |
| 12/1 | binder | ~200 nM | 0.870 | 0.880 | 0.007 | [d] |
| p53(17-29) | binder | ~580 nM | 0.830 | 0.840 | 0.007 | [d] |
| W23A | non-binder | >> 1 μM | 0.532 | 0.640 | 0.096 | [d] |
| poly-Ala | non-binder | — | 0.582 | 0.620 | 0.033 | [d] |
| scrambled | non-binder | — | 0.374 | 0.410 | 0.031 | [d] |

### 3.3 Expanded benchmark with hard negatives (Fig. S2)

The 29 hard negatives separated cleanly from the 3 known binders (Table 3). Minimum gap: +0.060 (lowest binder 0.850 vs highest hard negative 0.790). Cohen’s d = 3.41 [e] (95% CI: 1.94–4.88; Hedges’ g = 3.32; computed on individual iPTM values for all 29 negatives; full derivation in Supplementary Table S7; per-seed data in Supplementary Table S3), zero overlap. The W-position series revealed a distance-dependent gradient: W near the native position 8 (positions 10, 3, 1) retained partial signal (iPTM 0.68–0.79), while W at distal positions (2, 5, 11, 12) collapsed to non-binder range (0.37–0.48). Removal of Trp entirely (three no-W categories) converged on mean iPTM ∼0.43, consistent with Trp being the primary discriminant.

**Table 3.** Expanded AF3 benchmark: group summary (best iPTM, 5 seeds each). All data from [e].

| Group | N | Mean iPTM | SD | Range |
| --- | --- | --- | --- | --- |
| Known binders | 3 | 0.877 | 0.025 | 0.850–0.900 |
| W-position variants | 8 | 0.578 | 0.168 | 0.370–0.790 |
| Scrambled no-W | 5 | 0.438 | 0.062 | 0.380–0.530 |
| Hydrophobic no-W | 8 | 0.434 | 0.080 | 0.350–0.610 |
| Proteome-random | 8 | 0.432 | 0.077 | 0.290–0.550 |

### 3.4 Candidate screening

The 26 candidates spanned nine species. SPAFESTWDILK ranked 9th by AF3 iPTM (best 0.85, mean 0.83) and was selected for full evaluation based on: (i) 12-residue length matching PMI; (ii) derivation from ginger, an edible source; (iii) lowest 5 ns triage RMSD (0.13 nm, 2–5 ns window) [n]. This selection involved deliberate judgment beyond the AF3 score. Notably, SPAFESTWDILK ranked only #890/2,000 by ESM-2 similarity and would have been missed without the pharmacophore filter — the pipeline stages function as complementary, not redundant, filters.

### 3.5 Multi-method evaluation of SPAFESTWDILK

#### Three-platform co-folding concordance (Fig. S4)

Protenix yielded iPTM 0.923 (σ = 0.0003, 5 models [f], single-cluster pose convergence, max RMSD 0.37 Å). Chai-1 yielded iPTM 0.891 (σ = 0.001) [g]. Three-platform agreement (AF3 0.83 ± 0.01 mean / 0.85 best, Protenix 0.923, Chai-1 0.891) addresses single-architecture bias, with the caveat that all three tools share AlphaFold-derived architecture and are not fully independent (see Limitation 6). iPTM and pLDDT are confidence metrics, not binding affinity proxies; the ∼0.07 iPTM difference between AF3 and Protenix reflects implementation sensitivity. The predicted binding pose across all three platforms shows consistent three-pocket occupancy (Fig. 1).

**Fig. 1.**
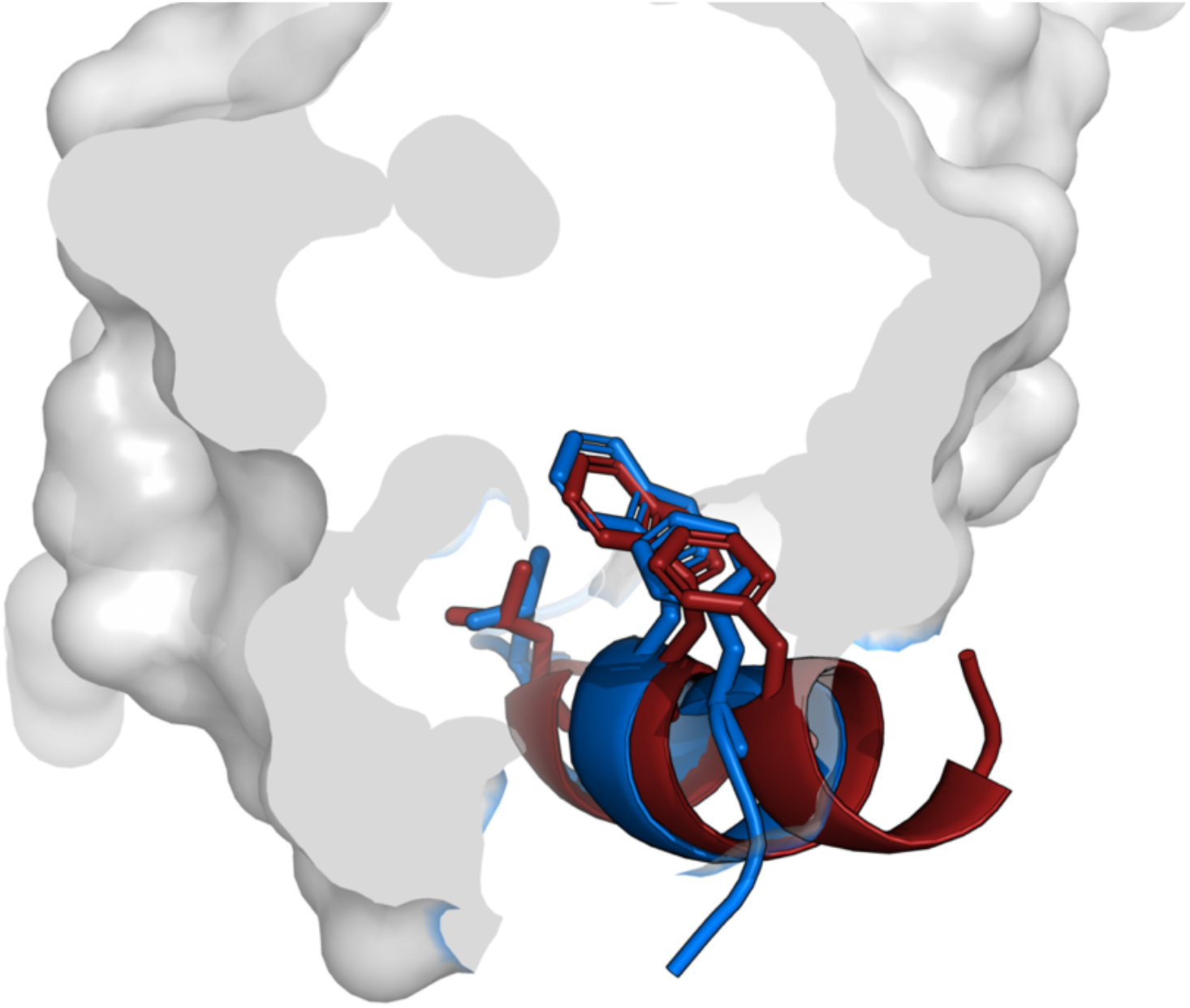
Structural mimicry of the p53 transactivation helix by SPAFESTWDILK in the MDM2 binding groove. Superimposition of the p53–MDM2 co-crystal structure (PDB 1YCR [1]; p53(17-29) in marine blue) with the AlphaFold 3 Server-predicted SPAFESTWDILK:MDM2 complex (SPAFESTWDILK in firebrick red), aligned on MDM2 Cα atoms (RMSD < 0.5 Å). Pharmacophore anchors shown as sticks: Phe4/Phe19 (green, Phe19 pocket), Trp8/Trp23 (blue, Trp23 pocket), Ile10–Leu11/Leu26 (red, Leu26 pocket). MDM2 surface shown in grey (30% transparency). Three-pocket occupancy is conserved across all three co-folding platforms (Fig. S4). Visualisation: open-source PyMOL. Scripts deposited at Zenodo DOI: 10.5281/zenodo.19859570 [x].

#### EvoEF2 (Supplementary Table S4)

Binding energy: −55.57 EEU (Protenix structure) [h], dominated by van der Waals (−39.16) and hydrophobic desolvation (−28.09). The AF3-based structure yielded −47.27 EEU [i]; this 8 EEU discrepancy reflects structural microstate sensitivity of the energy function, not a genuine energetic difference. Both values are separated from non-binder controls (scrambled −33.17, W23A −30.19 EEU). EEU = EvoEF2 Energy Units, not directly comparable to kcal/mol.

#### Molecular dynamics (**Fig. 2**)

Three independent Neurosnap simulations (AMBER99SB-ILDN 50 ns on Protenix structure [j], AMBER99SB-ILDN 50 ns on AF3 structure [k], CHARMM27 50 ns on AF3 structure [l]) plus a separate CHARMM36m [38] 100 ns run on Google Colab (performed on Google Colab A100; trajectory deposited at Zenodo DOI: 10.5281/zenodo.19859570) showed zero dissociation across two force fields and two starting structures. The CHARMM36m 100 ns run yielded RMSD 0.24 ± 0.05 nm (30–100 ns window; full trajectory 0.21 nm) [n] with pharmacophore core (FESTW) RMSF < 0.15 nm. The AMBER run showed RMSD 0.3–0.5 nm with transient excursions to ∼2 nm at 20–30 ns, representing conformational sampling of terminal residues while the core remained anchored. This cross-force-field variation is within the range documented for the p53-MDM2 system [14,19].

**Fig. 2.**
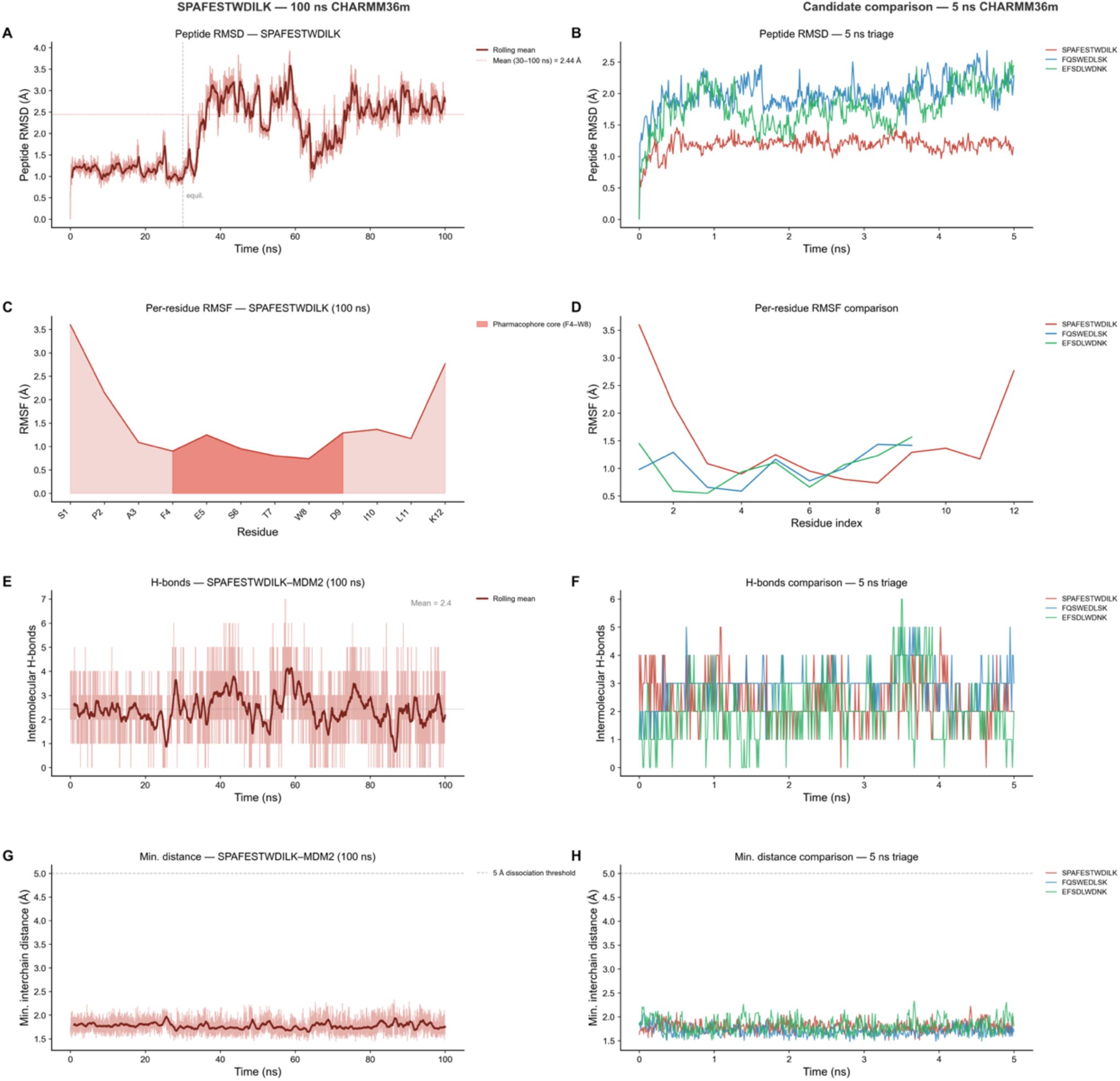
Molecular dynamics of the SPAFESTWDILK:MDM2 complex — 100 ns production run and 5 ns candidate triage. Left column: SPAFESTWDILK:MDM2 simulated for 100 ns under CHARMM36m force field (310 K, 0.15 M NaCl, TIP3P water; Google Colab A100 GPU, GROMACS 2023.1). Right column: 5 ns CHARMM36m triage comparing the three candidates promoted to extended simulation — SPAFESTWDILK (red), FQSWEDLSK (blue), and EFSDLWDNK (green). (A) Peptide backbone RMSD over 100 ns; dark red line = 50-frame rolling mean; dashed vertical line marks the 30 ns equilibration boundary; dotted horizontal line = equilibrated mean 2.44 Å (30–100 ns window). (B) Peptide RMSD comparison across three candidates at 5 ns; SPAFESTWDILK shows consistently lower RMSD than FQSWEDLSK and EFSDLWDNK, supporting its selection as lead. (C) Per-residue RMSF over 100 ns; shaded red region highlights the pharmacophore core F4–W8, which shows the lowest RMSF (<1.5 Å) confirming positional immobilisation within the binding groove. (D) Per-residue RMSF comparison across the three candidates; SPAFESTWDILK pharmacophore core is most rigid. (E) Intermolecular H-bonds (SPAFESTWDILK–MDM2 contacts only) over 100 ns; dark red = 100-frame rolling mean; mean = 2.4 bonds; dashed line = trajectory mean. (F) H-bond count comparison at 5 ns; all three candidates maintain comparable intermolecular contacts throughout the triage window. (G) Minimum interchain distance over 100 ns; dashed line at 5.0 Å = dissociation threshold; distance remains well below threshold throughout, confirming stable complex maintenance with no dissociation events. (H) Minimum distance comparison at 5 ns; all three candidates remain bound throughout the triage simulation. RMSD and RMSF in Å. Trajectory deposited at Zenodo DOI: 10.5281/zenodo.19859570. Raw XVG files: GROMACS_xvg_110files.zip [n].

#### MM-PBSA (**Fig. 3**; Supplementary Table S5)

AMBER yielded ΔG = −75.30 ± 4.92 kcal/mol [o]; CHARMM27 yielded −55.07 ± 2.86 kcal/mol [p]. The ∼20 kcal/mol difference reflects known electrostatic parameterization sensitivity (ΔEEL: −229 kcal/mol AMBER vs −86 CHARMM27). Both values are clearly negative and stable across the full sampling window (SEM 0.10 and 0.06 respectively; error bars show SEM across trajectory frames; total ΔG uncertainties reported as SD). These values exclude configurational entropy (−TΔS), which for a flexible 12-mer peptide can range from 10–30 kcal/mol [15]. Furthermore, the reported absolute values (−75.30 and −55.07 kcal/mol) are 4.7–6.5× larger in magnitude than the experimental ΔG of PMI (∼−11.6 kcal/mol at Kd = 3.3 nM), consistent with known MM-PBSA overestimation for charged peptide–protein systems [31]. These values support relative comparison between force fields and against controls but should not be interpreted as absolute binding free energies.

**Fig. 3.**
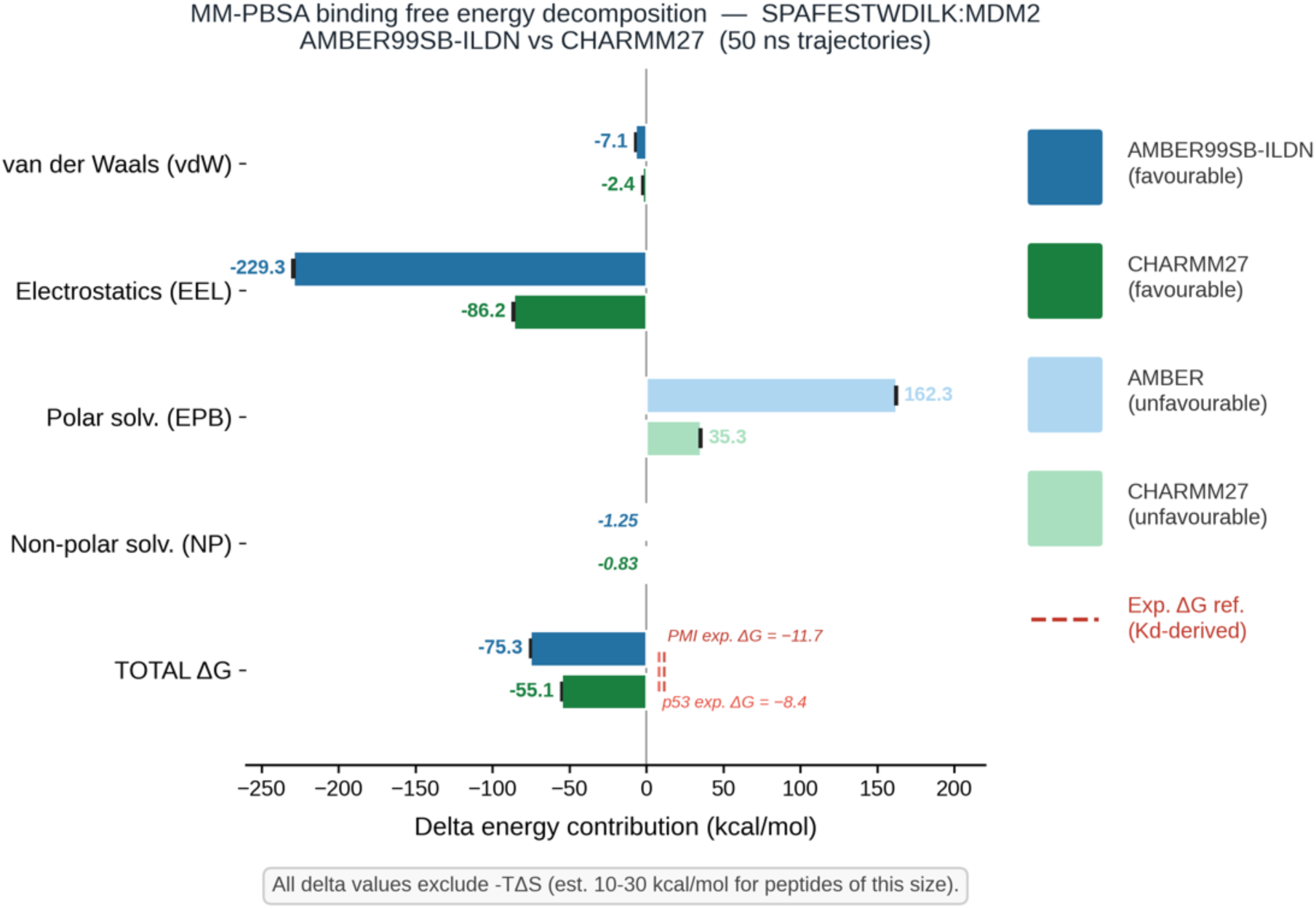
MM-PBSA binding free energy decomposition for the SPAFESTWDILK:MDM2 complex across two independent force fields. Horizontal grouped bars show MM-PBSA energy contributions (kcal/mol) for each component calculated from 50 ns GROMACS trajectories under AMBER99SB-ILDN (blue, Protenix structure [o]) and CHARMM27 (green, AF3 structure [p]). Favourable (negative) components are shown in dark blue/green; the unfavourable polar solvation penalty (ΔEPB) is shown in light blue/green. Error bars show SEM across trajectory frames; total ΔG uncertainties are reported as SD. ΔENPOLAR values are labelled directly on the plot (AMBER: −1.25; CHARMM27: −0.83 kcal/mol) as bars are not visible at this scale. Total ΔG: AMBER −75.30 ± 4.92 kcal/mol; CHARMM27 −55.07 ± 2.86 kcal/mol. Red dashed lines = experimental ΔG for PMI (−11.7 kcal/mol, Kd 3.3 nM) and p53(17-29) (−8.4 kcal/mol, Kd ∼580 nM), derived from Kd via ΔG = RT ln(Kd). The ∼20 kcal/mol inter-force-field difference is driven primarily by electrostatics, consistent with known differences in charge parameterisation between AMBER and CHARMM force fields. The 4–6× overestimation relative to experimental ΔG is expected in the absence of entropic correction (−TΔS, estimated 10–30 kcal/mol for peptides of this size [15,46]) and is consistent with known systematic overestimation of implicit-solvent MM-PBSA. Full component breakdown in Supplementary Table S5. Raw data at Zenodo DOI: 10.5281/zenodo.19859570.

### 3.6 Discrimination panel and W8A prediction

The seven-peptide panel on Protenix (Table 4) yielded clean separation. Cohen’s d = 6.4 (n = 7; df-weighted pooled SD; 95% CI: 2.75–10.12; Hedges’ g = 5.42; with n = 3 vs n = 4 this is a descriptive measure only), zero overlap. The W8A mutation caused an iPTM drop of 0.201 (0.923 to 0.722) [f,u], closely paralleling the W23A drop of 0.193 (0.920 to 0.727) (Fig. S3). W23A abolishes detectable p53-MDM2 binding [16]; in the PMI peptide context, the equivalent W23A mutation reduces affinity by > 50,000-fold (Kd from 3.3 nM to > 165 μM [2]). We predict that W8A substitution in SPAFESTWDILK will produce comparable binding loss. This constitutes the study’s principal testable hypothesis. The data support this prediction but do not establish biochemical equivalence between Trp8 and Trp23; confirmation requires SPR or ITC with the W8A mutant peptide.

**Table 4.** Discrimination panel (Protenix).

| Peptide | iPTM | Class | Ref |
| --- | --- | --- | --- |
| PMI (3.3 nM) | 0.940 | binder | [q] |
| SPAFESTWDILK | 0.923 | binder | [f] |
| p53(15-29) (140 nM) | 0.920 | binder | [r] |
| poly-Ala | 0.795 | non-binder | [s] |
| p53 W23A | 0.727 | non-binder | [t] |
| SPAFESTWDILK W8A | 0.722 | non-binder | [u] |
| scrambled | 0.718 | non-binder | [v] |

### 3.7 Peptide characterization

SPAFESTWDILK (MW 1,393.54 Da, pI 4.37, GRAVY −0.233, Boman index 1.24 kcal/mol [a,z]) corresponds to residues 344–355 of A0A8J5FLH2 [UniProt A0A8J5FLH2] (*Zingiber officinale* histone deacetylase, 731 aa, PE = 3). The peptide lies outside the HDAC catalytic domain (32–332), flanked by R343 and K355 (canonical tryptic sites). SPAFESTWDILK has not been reported in published ginger peptide inventories [7]; targeted proteomics (SRM/PRM) would be required to confirm its release from ginger hydrolysates. The F4-E5-S6-T7-W8 motif corresponds to the p53 F19-S20-D21-L22-W23 spacing; in the AF3 model, F4 occupies the Phe19 pocket, W8 the Trp23 pocket, and I10-L11 jointly the Leu26 pocket (Fig. S5). Residue contact network analysis confirms that SPAFESTWDILK shares all 16 MDM2 contacts with p53(17-29) with zero unique contacts (Fig. S6; Supplementary Table S6 for GI stability analysis).

## 4. DISCUSSION

### 4.1 Multi-method concordance

Eight computational assessments consistently classify SPAFESTWDILK as compatible with MDM2 binding: three co-folding architectures (AF3, Protenix, Chai-1 — all sharing AlphaFold-derived lineage), one knowledge-based energy function, two MD simulations with different force fields, and two MM-PBSA calculations. This concordance reflects robustness of a single binding hypothesis under diverse computational approximations, not independent thermodynamic validation, since the structural inputs share a common origin and the co-folding tools are not architecturally independent. Confidence metrics place SPAFESTWDILK between PMI (Kd = 3.3 nM) and p53(15-29) (Kd = 140 nM); these are model confidence rankings, not affinity measurements.

The cascaded pipeline architecture — combining proteome-scale *in silico* hydrolysis, physicochemical filtering, ESM-2 embedding-based dimensionality reduction, pharmacophore-driven selection, and multi-platform structural evaluation — extends and integrates approaches used in published multi-stage food-derived bioactive peptide workflows [27,28,41,42,43]. The integration of a protein language model embedding step as an intermediate dimensionality reduction layer, prior to pharmacophore filtering and structure-based scoring, has not been previously reported for a PPI cancer target. The observation that SPAFESTWDILK ranked only #890/2,000 by ESM-2 cosine similarity demonstrates that pipeline stages function as complementary, not redundant, filters.

SPAFESTWDILK shows lower conformational variability than p53(15-29) under identical simulation conditions (RMSD 0.24 vs ∼0.2 nm in published simulations [14]). This should not be interpreted as evidence of tighter binding: p53 is intrinsically disordered and its higher conformational flexibility in the MDM2 cleft is an intrinsic property, not a sign of weaker binding. The experimental Kd of p53(15-29) is 140 nM and RMSD comparisons cannot supersede this.

### 4.2 W8A as a testable prediction

The near-identical iPTM drops produced by W8A (Δ = 0.201) and W23A (Δ = 0.193), computed independently under identical conditions, support the prediction that Trp8 anchors SPAFESTWDILK into the MDM2 Trp23 sub-pocket analogously to Trp23 of p53. This is a falsifiable computational prediction, not a demonstrated biochemical equivalence. We recommend synthesis of SPAFESTWDILK and the W8A variant followed by SPR or ITC binding assay against recombinant MDM2(25–109) as the highest-priority experiment.

### 4.3 Limitations

The precursor protein A0A8J5FLH2 has PE = 3 status (inferred from homology, not experimentally detected in any *Z. officinale* tissue). The food-derived claim is contingent on experimental confirmation of protein expression in rhizome tissue; until such confirmation is obtained, SPAFESTWDILK should be considered a plant-database-derived computational candidate. AF3 thresholds were initially derived from n = 6; the expanded 36-peptide benchmark (Cohen’s d = 3.41, 95% CI: 1.94–4.88) mitigates but does not eliminate calibration uncertainty. MM-PBSA values exclude −TΔS (estimated 10–30 kcal/mol for peptides of this size [15]) and are 4.7–6.5× larger in magnitude than the experimental ΔG of PMI (∼−11.6 kcal/mol at Kd = 3.3 nM), consistent with known MM-PBSA overestimation for charged peptide–protein systems [35]; these values support relative comparison between force fields and against controls but should not be interpreted as absolute binding free energies. Sequential physicochemical filters bias the candidate pool toward canonical PPI chemotypes; non-aromatic or non-classical binders are systematically excluded. iPTM and pLDDT are confidence metrics, not thermodynamic quantities. The three co-folding tools (AF3, Protenix, Chai-1) share architectural lineage derived from AlphaFold and are not fully independent predictors; their agreement may partly reflect shared biases rather than independent confirmation. SPAFESTWDILK is predicted to be vulnerable to gastric and pancreatic proteases; oral delivery without stabilization is unlikely to yield bioavailable intact peptide (Supplementary Table S6).

### 4.4 Experimental roadmap

Priority experiments: (1) synthesis of SPAFESTWDILK and W8A variant; (2) SPR/BLI binding assay against MDM2(25–109) [gate-keeper experiment — if no binding detected, downstream steps are moot]; (3) ITC for direct thermodynamic measurement; (4) fluorescence polarization displacement assay; (5) simulated gastrointestinal stability and Caco-2 transport (only if adequate stability demonstrated); (6) LC-MS/MS identification in ginger tryptic hydrolysates; (7) cellular assays in p53-wild-type vs p53-null lines.

## 5. CONCLUSIONS

We report the computational identification and multi-method evaluation of SPAFESTWDILK, a dodecapeptide from *Zingiber officinale* histone deacetylase, as a predicted MDM2 binder. Eight computational assessments consistently classify the peptide as compatible with binding, with confidence metrics placing it between PMI (Kd = 3.3 nM) and p53(15-29) (Kd = 140 nM). A 36-peptide benchmark with hard negatives (composition- and filter-matched) supports the discriminative power of AF3 on this target (Cohen’s d = 3.41; 95% CI: 1.94–4.88; Hedges’ g = 3.32). The W8A/W23A iPTM parallel (Δ = 0.201 vs 0.193) constitutes the study’s principal testable prediction: W8A substitution should abolish SPAFESTWDILK binding to MDM2, analogous to the experimentally established W23A effect in p53. To our knowledge, this is the first food-database-derived linear peptide reported with multi-convergent computational evidence supporting engagement of the canonical three-anchor MDM2–p53 interface. Experimental validation is required.

## ACKNOWLEDGEMENTS

**General:** Computational analyses were performed on the Neurosnap platform, Google Colab, and Sherman Tree infrastructure. We acknowledge the open-source developers of GROMACS, gmx_MMPBSA, EvoEF2, and ESM-2. Statistical analysis and data science support were provided by StatExcel (https://statexcel.com). During the preparation of this work, the authors used AI-assisted writing tools for manuscript drafting and editing. After using these tools, the authors reviewed and edited all content as needed and take full responsibility for the content of the publication.

## Author contributions

Massimiliano Romiti: Conceptualization, Methodology, Software, Investigation, Resources, Writing – Original Draft, Project administration, Funding acquisition, Supervision. Minoo Ashtiani: Conceptualization, Methodology, Software, Validation, Formal analysis, Visualization, Writing – Review & Editing. Carla Sandri: Software, Validation, Writing – Review & Editing. Giulia Paiola: Validation, Writing – Review & Editing.

## Data availability

All raw computational outputs are deposited at Zenodo DOI: 10.5281/zenodo.19859570 (paper outputs) and Zenodo DOI: 10.5281/zenodo.20447996 (proteome database). Pipeline code (PepVeg screening notebook, single-filter mode notebook, and GROMACS CHARMM36m setup notebook) is available at https://github.com/Kingelanci/PepVeg-Pipeline. Neurosnap platform job links for all independent multi-method evaluation analyses are listed in Supplementary Table S1.

