## Supplementary Figures and Tables for "SPAFESTWDILK, a plant-derived dodecapeptide from *Zingiber officinale*, as a predicted inhibitor of the MDM2–p53 interaction: computational discovery and multi-method evaluation" (from another version of this preprint: version 1, 2026-06-10)

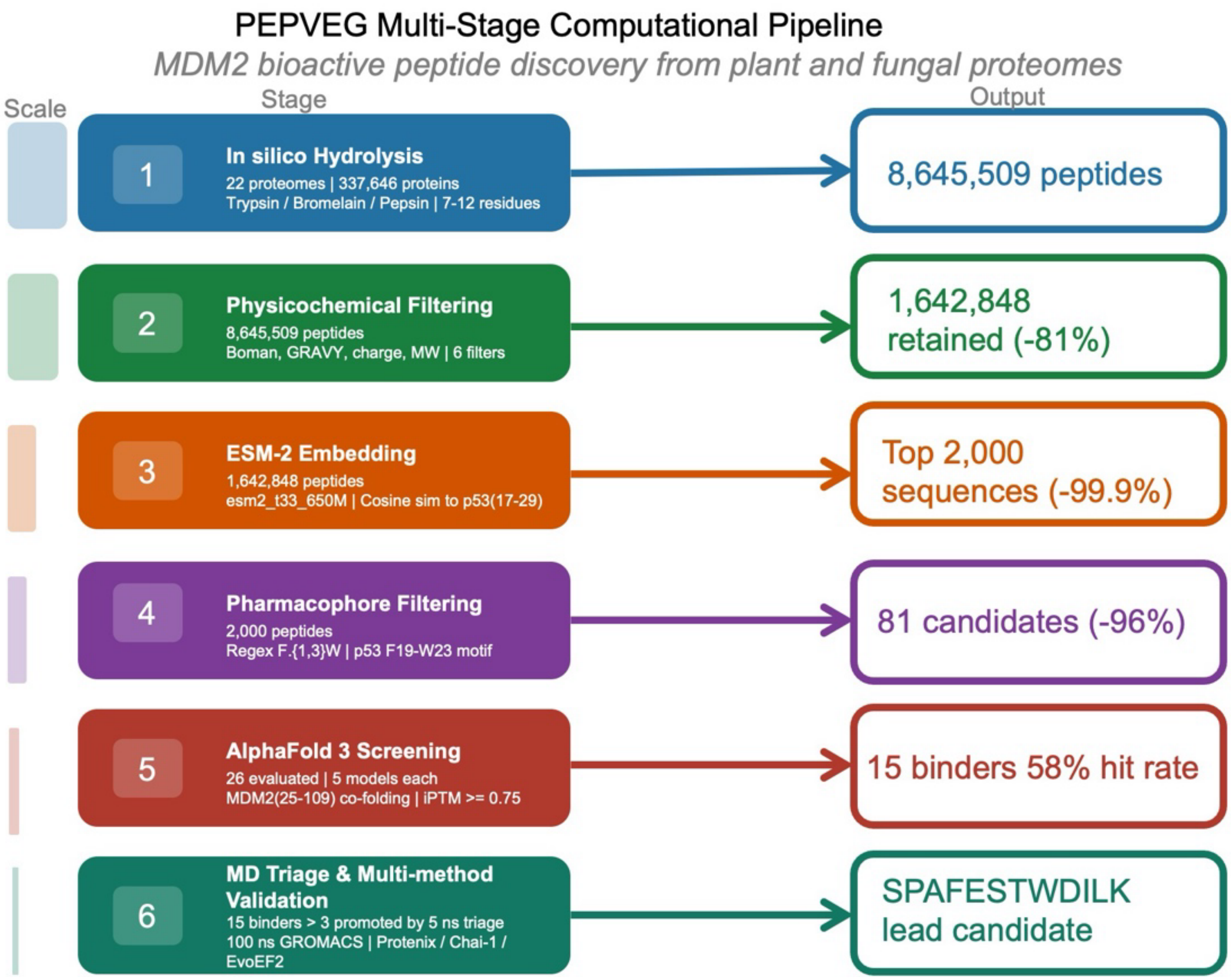


**Fig. S1. PEPVEG multi-stage computational pipeline for MDM2 bioactive peptide discovery from plant and fungal proteomes.**

Six-stage pipeline schematic applied to 22 plant and fungal proteomes (337,646 proteins). Stage 1 (blue): in silico tryptic, bromelain, and pepsin hydrolysis yields 8,645,509 unique 7–12 residue fragments. Stage 2 (green): six sequential physicochemical filters (Boman index ≥1.0, GRAVY −2.0 to +0.5, net charge −2 to +4, instability index <40, MW <1,500 Da, hydrophobic fraction 30–60%) reduce the pool to 1,642,848 peptides (−81.0%). Stage 3 (orange): ESM-2 embedding (esm2_t33_650M_UR50D) and cosine similarity ranking against p53(17–29) retains the top 2,000 sequences (−99.9%). Stage 4 (purple): pharmacophore regex F.{1,3}W (encoding the p53 F19–W23 spacing) selects 81 candidates (−96.0%). Stage 5 (dark red): AlphaFold 3 co-folding of 26 selected candidates against MDM2(25–109) identifies 15 putative binders (iPTM ≥0.75; 58% hit rate). Stage 6 (teal): 5 ns GROMACS CHARMM36m triage followed by 100 ns production simulation and multi-method validation (Protenix, Chai-1, EvoEF2, MM-PBSA) yields SPAFESTWDILK as the lead candidate. Left-hand funnel bars indicate the relative scale of each output on a compressed logarithmic scale.

*SPAFESTWDILK ranked #890/2,000 by ESM-2 cosine similarity and was recovered exclusively through pharmacophore filtering (Stage 4). ESM-2 cosine similarity shows near-zero correlation with AF3 iPTM (Pearson r = −0.049, n = 97 control peptides), establishing the complementary and non-redundant nature of pipeline stages. Filter parameters in Supplementary Table S2.*

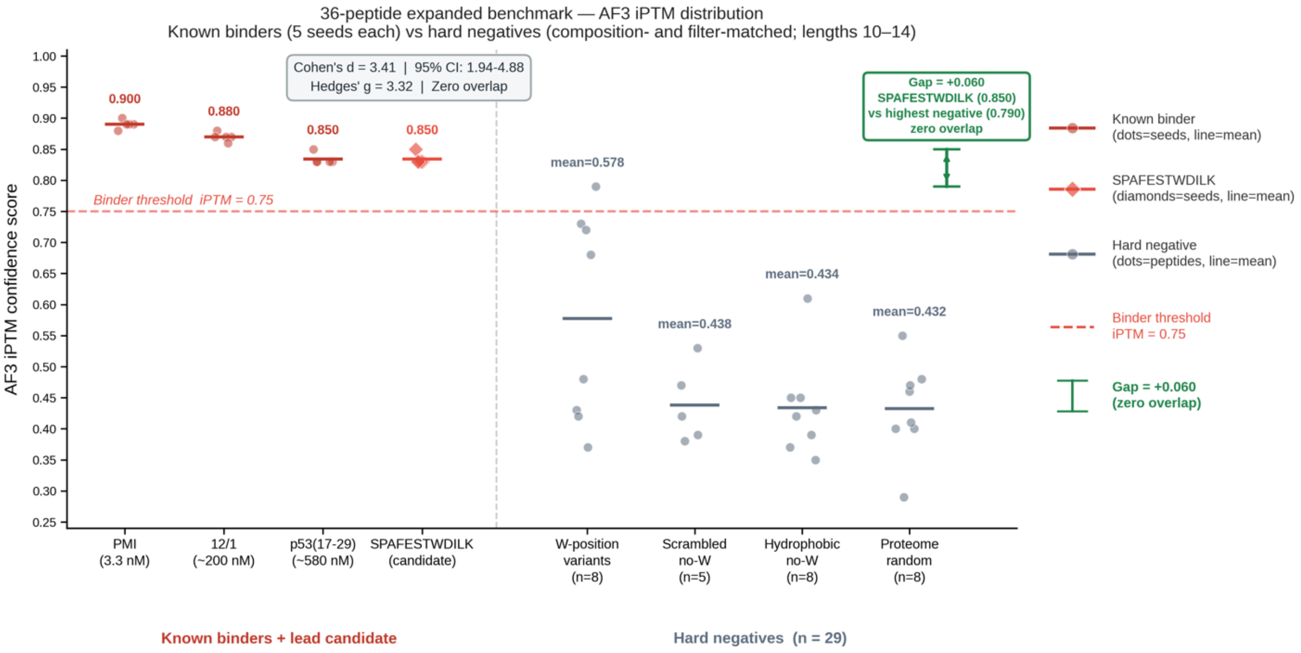


**Fig. S2. 36-peptide expanded AF3 benchmark: iPTM distribution for known binders and hard negatives.**

AF3 Server best-seed iPTM scores for three reference binders (filled symbols, individual values with Kd annotated), the lead candidate SPAFESTWDILK (filled diamond, iPTM 0.850), and 29 hard negatives in four categories (mean ± full range). Hard negative categories: W-position variants (n = 8, mean 0.578, range 0.370–0.790) — Trp relocated from position 8 to non-native positions, lengths 11–12; Scrambled no-W (n = 5, mean 0.438, range 0.380–0.530) — W8A substitution with full sequence scrambling, length 12; Hydrophobic no-W (n = 8, mean 0.434, range 0.350–0.610) — W8 replaced by a hydrophobic residue (Leu/Ile/Val/Met/Phe), lengths 11–12; Proteome-random (n = 8, mean 0.432, range 0.290–0.550) — tryptic peptides from housekeeping proteins passing the same six physicochemical filters as SPAFESTWDILK, lengths 10–14. The first three categories are composition-matched to SPAFESTWDILK; the proteome-random category is filter-matched only. Length was not strictly controlled (range 10–14 vs SPAFESTWDILK 12 residues). Zero overlap between all binder values (minimum 0.850) and all negative values (maximum 0.790); gap = +0.060. Dashed line: binder threshold iPTM = 0.75. Effect size: Cohen's d = 3.41 (95% CI: 1.94–4.88); Hedges' g = 3.32.

*All evaluations on AF3 Server (alphafoldserver.com), 5 seeds per peptide, MDM2(25–109) construct. Per-seed data in Supplementary Table S3. Effect size derivation in Supplementary Table S7.*

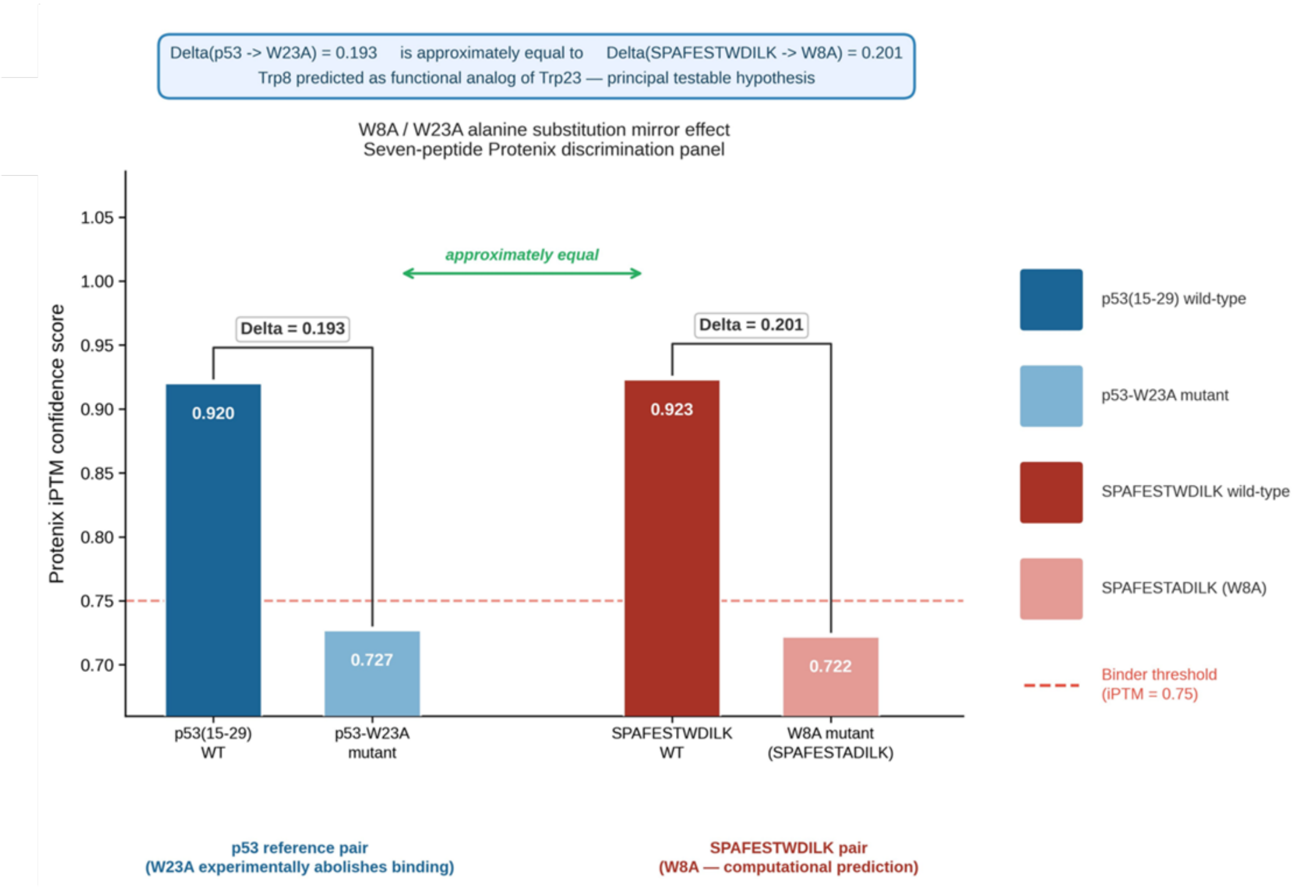


**Fig. S3. W8A/W23A alanine substitution mirror effect: Trp8 as a functional analogue of Trp23.**

Protenix iPTM scores for wild-type and Ala-substituted pairs from the seven-peptide discrimination panel (Neurosnap platform, MDM2(25–109), 5 models each). Left pair: p53(15–29) wild-type (iPTM 0.920) vs p53-W23A (iPTM 0.727), Δ = 0.193. W23A abolishes detectable p53–MDM2 binding in vitro and reduces PMI affinity by >50,000-fold (Pazgier et al. 2009). Right pair: SPAFESTWDILK (iPTM 0.923) vs SPAFESTADILK W8A (iPTM 0.722), Δ = 0.201. The near-identical Δ values (difference = 0.008) support the prediction that Trp8 occupies the MDM2 Trp23 sub-pocket analogously to p53 Trp23. W8A substitution is predicted to abolish SPAFESTWDILK binding — the study's principal falsifiable experimental prediction. Dashed line: binder threshold iPTM = 0.75.

*Protenix job IDs in Supplementary Table S1. Full 7-peptide panel effect size: Cohen's d = 6.43 (Hedges' g = 5.42), derived in Supplementary Table S7.*

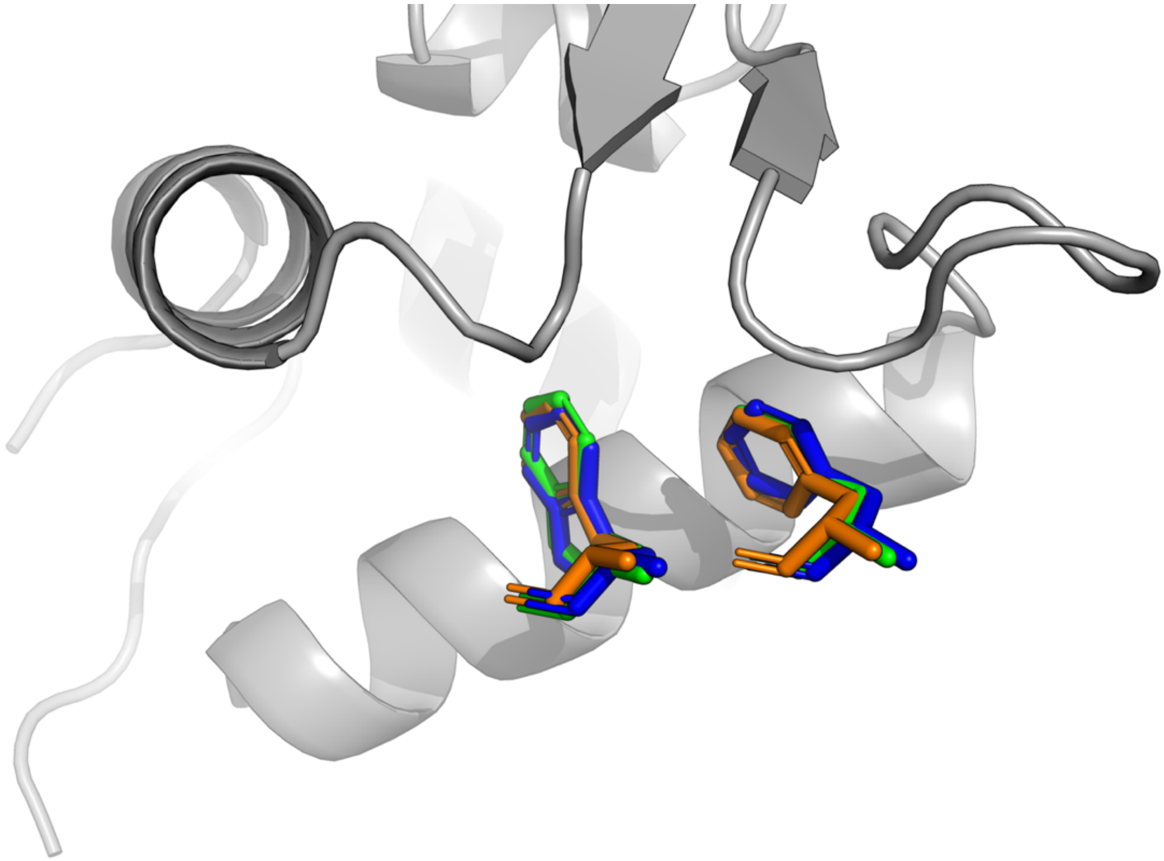

**Fig. S4. Cross-platform pose convergence for SPAFESTWDILK:MDM2 across three co-folding architectures.**

Superimposition of predicted SPAFESTWDILK:MDM2 binding poses from AlphaFold 3 Server (marine blue, best iPTM 0.850), Protenix (forest green, iPTM 0.923), and Chai-1 (orange, iPTM 0.891), aligned on MDM2 chain A Cα atoms. MDM2 N-terminal domain shown as grey cartoon. Stick representations show Trp8 indole and Phe4 phenyl sidechain positions across all three models; the three indole rings overlay within a maximum pairwise RMSD of 0.37 Å (Protenix, 5 models), and Phe4 rings show equivalent convergence, demonstrating single-cluster pose agreement across independently implemented co-folding platforms. Caveat: AF3, Protenix, and Chai-1 share AlphaFold-derived architectural lineage; convergence may partly reflect shared training biases rather than fully orthogonal validation.

*Visualisation: open-source PyMOL. AF3: alphafoldserver.com (5 seeds). Protenix and Chai-1: Neurosnap platform (5 models each). Neurosnap job IDs in Supplementary Table S1.*

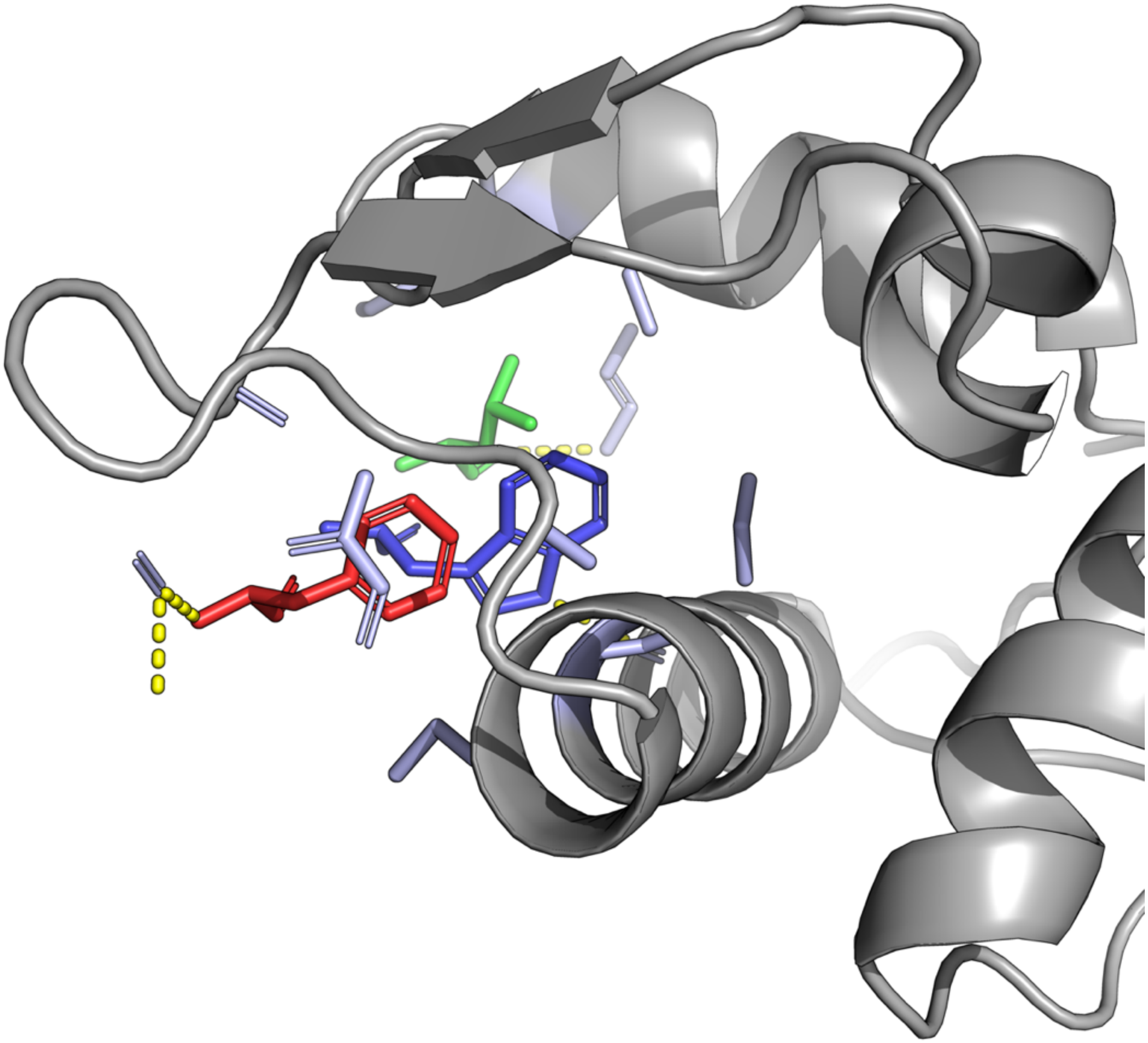

**Fig. S5. SPAFESTWDILK:MDM2 binding site contacts in the AlphaFold 3 predicted complex.**

Three-dimensional representation of intermolecular contacts between SPAFESTWDILK (coloured sticks) and the MDM2 N-terminal domain (grey cartoon) in the AF3 Server top-ranked structure (MDM2 residues 25–109). Pharmacophore anchor sidechains are colour-coded by sub-pocket: Phe4 (green, Phe19 sub-pocket), Trp8 (blue, Trp23 sub-pocket — deepest cavity and primary anchor), and Leu11 (red, Leu26 sub-pocket). MDM2 contact residues within 4.5 Å of each anchor sidechain are shown as light blue sticks. Yellow dashed lines indicate predicted hydrogen bonds (distance ≤3.2 Å). The Trp8 indole ring inserts deepest into the groove, consistent with its role as the primary pharmacophore anchor confirmed by the W8A iPTM drop of 0.201 (Figure S3).

*Visualisation: open-source PyMOL. Input: AF3 Server top-ranked model, run fold_2026_04_14_09_55 (chain B residues 1–12, chain A residues 25–109). Contact cutoff: 4.5 Å. H-bond cutoff: 3.2 Å.*

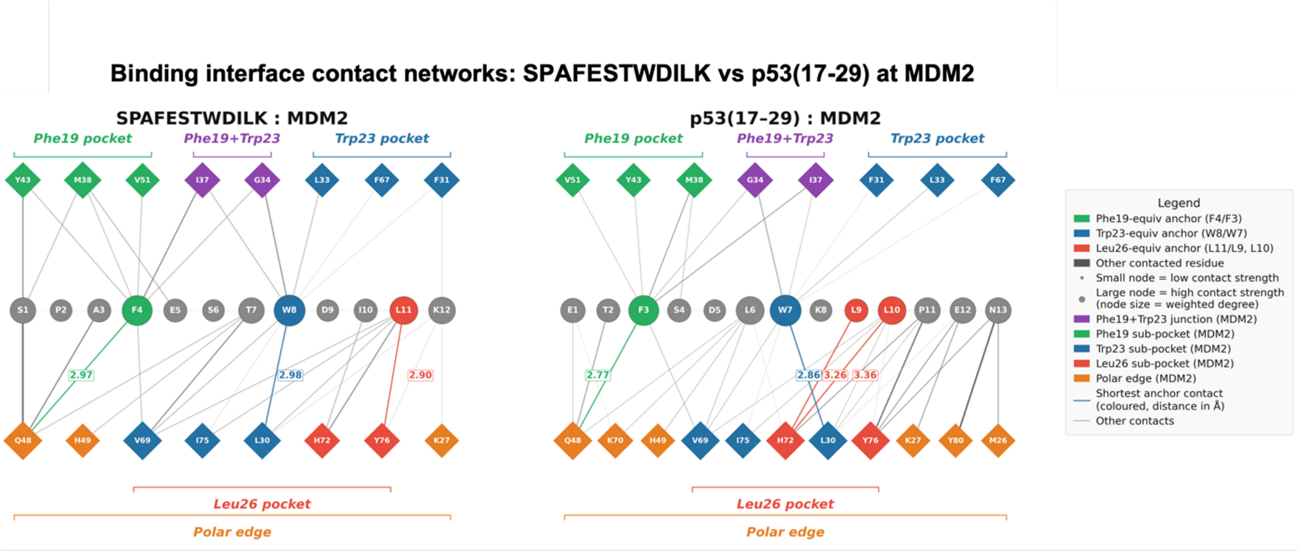


**Fig. S6. Residue contact network analysis of the SPAFESTWDILK:MDM2 and p53(17–29):MDM2 binding interfaces.**

Bipartite contact networks for SPAFESTWDILK:MDM2 (left) and p53(17–29):MDM2 (right), computed from AF3 Server top-ranked structures. Circular nodes represent peptide residues (left to right: N- to C-terminus); diamond nodes represent MDM2 residues. Edges connect residue pairs with minimum heavy-atom distance ≤5.0 Å. Edge thickness is proportional to contact strength (1/distance). Node size encodes normalised weighted degree (WD = $\Sigma(1/d_{ij})$, normalised to the maximum value in each network); larger nodes indicate higher total contact strength. Peptide pharmacophore anchors are colour-coded: Phe19-equivalent (F4/F3, green), Trp23-equivalent (W8/W7, blue), Leu26-equivalent (L11/L9, L10, red). MDM2 residues are coloured by sub-pocket: Phe19 (green diamonds), Trp23 (blue), Leu26 (red), polar edge (orange), Phe19+Trp23 junction G34/I37 (purple). Coloured highlighted edges show the shortest anchor contact with distance in Å. Sub-pocket zone brackets annotated above and below each panel. SPAFESTWDILK shares all 16 MDM2 contacts with p53(17–29) with zero unique contacts; the three p53-exclusive contacts (K70, M26, Y80) are made by p53 residues with no structural equivalent in SPAFESTWDILK. Weighted degree centrality identifies W8/W7 (WD = 1.00) as the dominant hub in both interfaces, followed by F4/F3 (WD = 0.92/0.99) and L11/L10 (WD = 0.67/0.61).

*Both structures from AF3 Server: SPAFESTWDILK run fold_2026_04_14_09_55; p53(17–29) = ETFSDLWKLLPEN (13 residues, Kd ~580 nM, Kussie et al. 1996) run fold_2026_04_13_13_03. Note: the longer p53(15–29) fragment (15 residues, Kd 140 nM) was used in the Protenix discrimination panel (Figure S3, Table S7); the structural comparison here uses p53(17–29) as in PDB 1YCR. Contact computation: minimum heavy-atom distance, 5.0 Å cutoff. Network analysis: Python, NetworkX. Raw coordinate files at Zenodo DOI: 10.5281/zenodo.19859570*

### SUPPLEMENTARY TABLES

#### Supplementary Table S1. Neurosnap platform job links.

Public Neurosnap job links for all computational analyses performed on the Neurosnap platform (neurosnap.ai), organised by analysis type: GROMACS molecular dynamics (four 50 ns simulations: AMBER99SB-ILDN Protenix structure, AMBER99SB-ILDN AF3 structure, CHARMM27 AF3 structure, and p53(17–29) reference control), co-folding and energy scoring (Protenix iPTM 0.923, Chai-1 iPTM 0.891, EvoEF2 −55.57 EEU [EvoEF2 Energy Units]), MM-PBSA binding free energy (AMBER −75.30 ± 4.92 kcal/mol; CHARMM27 −55.07 ± 2.86 kcal/mol), and the seven-peptide Protenix discrimination panel. Each entry includes category, description, platform, key parameters, and full public read-only share URL.

*AF3 Server jobs were performed directly on alphafoldserver.com and are not listed here; raw JSON outputs are deposited at Zenodo DOI: 10.5281/zenodo.19859570. The 100 ns CHARMM36m simulation was performed on Google Colab A100 GPU (GROMACS 2023.1); raw XVG files deposited at the same Zenodo repository.*

| Category | Description | Platform | Key parameters | Public URL |
|---|---|---|---|---|
| **GROMACS Molecular Dynamics Simulations** | | | | |
| **GROMACS — AMBER99SB-ILDN (Protenix struct)** | 50 ns MD simulation of SPAFESTWDILK:MDM2(25-109); Protenix rank_1 structure as input | Neurosnap | *AMBER99SB-ILDN, TIP3P, 310 K, 0.15 M NaCl, 50 ns* | neurosnap.ai/job/69df8c8dc7a74174eaf1a6d5?share=69e22151eec1918b94662216 |
| **GROMACS — AMBER99SB-ILDN (AF3 struct)** | 50 ns MD simulation of SPAFESTWDILK:MDM2(25-109); AF3 Server model_0 as input | Neurosnap | *AMBER99SB-ILDN, TIP3P, 310 K, 0.15 M NaCl, 50 ns* | neurosnap.ai/job/69e0bca1c7a74174eaf1b631?share=69e22151eec1918b9466221a |
| **GROMACS — CHARMM27 (AF3 struct)** | 50 ns MD simulation of SPAFESTWDILK:MDM2(25-109); AF3 Server model_0 as input | Neurosnap | *CHARMM27, TIP3P, 310 K, 0.15 M NaCl, 50 ns* | neurosnap.ai/job/69e0f5b3c7a74174eaf1b9d9?share=69e22151eec1918b9466221c |
| **GROMACS — p53(17-29) control (AMBER99SB-ILDN)** | 50 ns MD simulation of p53(17-29):MDM2(25-109); reference control | Neurosnap | *AMBER99SB-ILDN, TIP3P, 310 K, 0.15 M NaCl, 50 ns* | neurosnap.ai/job/69e12b80c7a74174eaf1bc65?share=69e22151eec1918b9466221e |
| **Co-folding and Energy Scoring** | | | | |
| **Protenix co-folding — SPAFESTWDILK:MDM2** | Cross-platform co-folding, best iPTM = 0.923; structure used for EvoEF2 and AMBER MD | Neurosnap | *MDM2(25-109), 5 models, SPAFESTWDILK* | neurosnap.ai/job/69df686cc7a74174eaf1a524?share=69e22150eec1918b946621fe |

| Category | Description | Platform | Key parameters | Public URL |
|---|---|---|---|---|
| **Chai-1 co-folding — SPAFESTWDILK:MDM2** | Cross-platform co-folding, best iPTM = 0.891 | Neurosnap | *MDM2(25-109), 5 models, SPAFESTWDILK* | neurosnap.ai/job/69df6680c7a74174eaf1a51c?share=69e22150eec1918b9466220e |
| **EvoEF2 — SPAFESTWDILK:MDM2 (Protenix struct)** | Knowledge-based energy decomposition on Protenix rank_1 structure; score = -55.57 EEU. AF3 model_0 score = -47.27 EEU (see Supplementary Table S4). | Neurosnap | *Protenix rank_1 input, chain A vs B* | neurosnap.ai/job/69df74bfc7a74174eaf1a585?share=69e22151eec1918b94662212 |
| **MM-PBSA Binding Free Energy** | | | | |
| **MM-PBSA — AMBER99SB-ILDN (50 ns)** | Binding free energy decomposition; DeltaG_total = -75.30 +/- 4.92 kcal/mol | Neurosnap | *AMBER99SB-ILDN trajectory, gmx_MMPBSA, implicit solvent* | neurosnap.ai/job/69dfc05fc7a74174eaf1a983?share=69e22152eec1918b94662222 |
| **MM-PBSA — CHARMM27 (50 ns)** | Binding free energy decomposition; DeltaG_total = -55.07 +/- 2.86 kcal/mol | Neurosnap | *CHARMM27 trajectory, gmx_MMPBSA, implicit solvent* | neurosnap.ai/job/69e12743c7a74174eaf1bc52?share=69e22152eec1918b94662224 |
| **7-Peptide Protenix Discrimination Panel** | | | | |
| **Protenix discrimination — SPAFESTWDILK (binder)** | 7-peptide panel; iPTM = 0.923; used in W8A/W23A mirror analysis | Neurosnap | *MDM2(25-109), 5 models* | neurosnap.ai/job/69df686cc7a74174eaf1a524?share=69e22150eec1918b946621fe |
| **Protenix discrimination — PMI (ctrl+)** | Positive control binder; Kd 3.3 nM; iPTM = 0.940 | Neurosnap | *MDM2(25-109), 5 models* | neurosnap.ai/job/69df89d5c7a74174eaf1a688?share=69e22150eec1918b94662202 |
| **Protenix discrimination — p53(15-29) (ctrl+)** | Positive control binder; Kd ~140 nM; iPTM = 0.920 | Neurosnap | *MDM2(25-109), 5 models* | neurosnap.ai/job/69df89bbc7a74174eaf1a686?share=69e22150eec1918b94662204 |
| **Protenix discrimination — Poly-Ala (ctrl-)** | Negative control non-binder; iPTM = 0.795 | Neurosnap | *MDM2(25-109), 5 models* | neurosnap.ai/job/69df8844c7a74174eaf1a678?share=69e22150eec1918b94662206 |
| **Protenix discrimination — p53 W23A (ctrl-)** | Negative control; W23A abolishes p53 binding; iPTM = 0.727 | Neurosnap | *MDM2(25-109), 5 models* | neurosnap.ai/job/69df89f1c7a74174eaf1a68a?share=69e22150eec1918b94662208 |
| **Protenix discrimination — SPAFESTADILK W8A (ctrl-)** | W8A substitution of lead candidate; iPTM = 0.722; principal falsifiable prediction | Neurosnap | *MDM2(25-109), 5 models* | neurosnap.ai/job/69df8956c7a74174eaf1a682?share=69e22150eec1918b9466220a |

| Category | Description | Platform | Key parameters | Public URL |
|---|---|---|---|---|
| **Protenix discrimination — Scrambled (ctrl-)** | Scrambled sequence negative control; iPTM = 0.718 | Neurosnap | *MDM2(25-109), 5 models* | neurosnap.ai/job/69df88acc7a74174eaf1a67f?share=69e22150eec1918b9466220c |

*Note: Neurosnap share links provide read-only public access to job inputs, parameters, outputs, and result files. URLs are stable permalinks. AF3 Server = alphafoldserver.com (5 seeds per peptide, MDM2 25-109 construct, no MSA). Protenix and Chai-1 = Neurosnap implementations (5 models each, MDM2 25-109 construct).*

### Supplementary Table S2. Physicochemical filter parameters.

Complete specification of the six sequential physicochemical filters applied in Stage 2 of the PEPVEG pipeline to 8,645,509 unique peptide fragments. For each filter: threshold value, formula with scale reference, scientific rationale, literature reference, and the verified SPAFESTWDILK value. Intermediate counts from single-filter mode pipeline rerun (RunPod, NutrAI Pipeline v9, May 2026; same environment as the original run): Boman index (5,127,842 passing independently, 59.31% — most restrictive), GRAVY (6,870,519, 79.47%), net charge (8,021,155, 92.78%), instability index (8,448,408, 97.72% — least restrictive), molecular weight (5,959,152, 68.93%), hydrophobic fraction (5,559,756, 64.31%). Combined six-filter sequential output: 1,642,848 peptides (−81.0%), consistent with the manuscript value. SPAFESTWDILK passes all six filters: Boman index 1.24, GRAVY −0.23, net charge −1 (E:−1, D:−1, K:+1), instability index 37.67, MW 1,393.5 Da, hydrophobic fraction 50% (P, A, F, W, I, L).

*Filters constitute chemical space reduction and introduce bias toward canonical PPI-targeting chemotypes; they are not biologically validated selection criteria. Implementation in VegPep_Pipeline_v10.ipynb (Zenodo DOI: 10.5281/zenodo.19859570).*

#### Filter specifications

| # | Filter | Threshold | Formula / Implementation | Scientific rationale | Reference | SPAFESTWDILK value |
|---|---|---|---|---|---|---|
| 1 | **Boman Index** | **>= 1.0** | *Boman = -( sum(DeltaG_transfer_i) ) / n DeltaG values: Radzicka & Wolfenden 1988 (kcal/mol, cyclohexane->water)* | Predicts protein-binding potential. Higher values indicate greater likelihood of | *Boman 2003; Radzicka & Wolfenden 1988* | **1.24 (PASS)** |

| # | Filter | Threshold | Formula / Implementation | Scientific rationale | Reference | SPAFESTWDILK value |
|---|---|---|---|---|---|---|
| | | | | protein-protein interaction involvement. Threshold >= 1.0 enriches for PPI-compatible chemotypes. | | |
| 2 | **GRAVY (Grand Average of Hydropathicity)** | **-2.0 to +0.5** | *GRAVY = sum(Kyte-Doolittle_i) / n Kyte-Doolittle scale (1982)* | Excludes highly hydrophobic peptides (membrane-targeting) and highly hydrophilic peptides (likely disordered). Range selects for amphipathic, PPI-compatible sequences. | *Kyte & Doolittle 1982* | **-0.23 (PASS)** |
| 3 | **Net charge** | **-2 to +4** | *Charge = sum(R:+1, K:+1, H:+0.5, D:-1, E:-1)* | MDM2 binding groove is predominantly hydrophobic; excludes highly charged peptides that would not penetrate the groove. Slight positive bias reflects p53 charge pattern. | *Based on p53(17-29) charge profile* | **-1 (PASS; E:-1, D:-1, K:+1)** |
| 4 | **Instability Index** | **< 40** | *II = (10/n) * sum(DIWV[seq[i]+seq[i+1]]) DIWV = dipeptide instability weight values (Guruprasad 1990)* | Predicts in vitro stability. Index < 40 classifies peptide as stable. Excludes sequences likely to degrade before experimental validation. | *Guruprasad et al. 1990* | **37.67 (PASS)**[1] |
| 5 | **Molecular weight** | **< 1,500 Da** | *MW = sum(MW_residue_i) - (n-1) * 18.02 Residue MW values: standard monoisotopic* | Upper limit for oral bioavailability consideration and typical PPI peptide size range. SPAFESTWDILK (12 residues) falls well within this range. | *Lipinski-inspired; empirical PPI peptide range* | **1,393.5 Da (PASS)** |
| 6 | **Hydrophobic fraction** | **30% to 60%** | *HF = count(A,I,L,M,F,P,W,V) / n * 100 Hydrophobic residues: AILMFPWV* | MDM2 Trp23/Phe19/Leu26 sub-pockets are | *Based on p53 pharmacophore anchor analysis* | **50% (PASS; P+A+F+W+I+L = 6/12)** |

[1] Pipeline implementation value (simplified DIWV matrix). ProtParam (complete 400-entry DIWV matrix) yields 74.33, which would not pass this filter; see Section 2.3 for full disclosure.

| # | Filter | Threshold | Formula / Implementation | Scientific rationale | Reference | SPAFESTWDILK value |
|---|---|---|---|---|---|---|
| | | | | hydrophobic. Too low hydrophobicity = poor groove fit; too high = aggregation / membrane insertion risk. | | |

### Pipeline funnel — Step 2 sequential filter outputs

| Stage | Input | Filter applied | Output | Reduction |
|---|---|---|---|---|
| **1 — Enzymatic hydrolysis** | 337,646 proteins (22 proteomes) | *Trypsin + Bromelain + Pepsin; 7-12 residues* | 8,645,509 peptides | - |
| **2a — Boman Index** | 8,645,509 | **>= 1.0** | **5,128,049** | −3,517,460 (−40.69%) |
| **2b — GRAVY** | 5,128,049 | **-2.0 to +0.5** | **4,573,283** | −554,766 (−6.42%) |
| **2c — Net charge** | 4,573,283 | **-2 to +4** | **4,091,058** | −482,225 (−5.58%) |
| **2d — Instability Index** | 4,091,058 | **< 40** | **3,988,394** | −102,664 (−1.19%) |
| **2e — Molecular weight** | 3,988,394 | **< 1,500 Da** | **2,864,569** | −1,123,825 (−13.00%) |
| **2f — Hydrophobic fraction** | 2,864,569 | **30–60%** | **1,642,848** | −1,221,721 (−14.13%) |
| **2 — Combined (all 6 filters)** | 8,645,509 | All 6 sequential | **1,642,848 peptides retained** | **−81.0%** |

*Intermediate counts from single-filter mode pipeline rerun (RunPod, NutrAI Pipeline v9, May 2026). All counts verified on the same RunPod environment used for the original pipeline run; the combined output of 1,642,848 is identical to the manuscript value. The dominant filter is Boman index (removes 40.69% of input independently); hydrophobic fraction and MW are second and third most restrictive. Instability index is the least restrictive filter (removes only 2.28%). Pipeline notebook: https://github.com/Kingelanci/PepVeg-Pipeline*

### Supplementary Table S3. Expanded 36-peptide AF3 benchmark — complete per-seed iPTM data.

Per-seed AlphaFold 3 Server iPTM values for 32 of the 36 peptides in the expanded benchmark: three reference binders (PMI, 12/1, SPAFESTWDILK) and 29 hard negatives across four categories (W-position variants n = 8, scrambled no-W n = 5, hydrophobic no-W n = 8, proteome-random n = 8). The remaining four peptides (p53(17–29), p53-W23A, poly-Ala, Scrambled) are AF3 Server controls listed in Supplementary Table S1; their Protenix iPTM values are in Supplementary Table S7 Panel B. All evaluations used AF3 Server (alphafoldserver.com), MDM2(25–109), no MSA for the peptide chain, five independent random seeds per peptide. Best iPTM, mean, and SD across five seeds are reported. Sequences verified from AF3 Server job_request JSON files; all 29 hard-negative best-iPTM values cross-validated against the Supplementary Table S7 derivation file. Matching criteria: W-position variants, scrambled no-W, and hydrophobic no-W categories were composition-matched to SPAFESTWDILK (same or analogous residues, lengths 11–12); proteome-random peptides were filter-matched only (same six physicochemical filters as Table S2; lengths 10–14). Length was not strictly controlled across categories.

*All evaluations on AF3 Server (alphafoldserver.com), MDM2(25–109) construct, no MSA for peptide chain. Raw JSON files in AF3_Server_Benchmark_29_Hard_Negatives.zip (Zenodo DOI: 10.5281/zenodo.19859570). Effect size statistics derived in Supplementary Table S7.*

*Shading key: Green = binders; Blue = W-position variants; Teal = scrambled no-W; Amber = hydrophobic no-W; Purple = proteome-random.*

| ID / Peptide | Sequence | Category | Kd | Seed 1 | Seed 2 | Seed 3 | Seed 4 | Seed 5 | Best iPTM | Mean iPTM | SD |
|---|---|---|---|---|---|---|---|---|---|---|---|
| **BINDERS — 3 known binders / lead candidate (AF3 Server, 5 seeds, MDM2(25-109))** | | | | | | | | | | | |
| **PMI** | TSFAEYWNLLSP | *Known binder* | 3.3 nM | 0.90 | 0.89 | 0.89 | 0.89 | 0.88 | **0.900** | 0.890 | 0.007 |
| **12/1** | MPRFMDYWEGLN | *Known binder* | ~200 nM | 0.88 | 0.87 | 0.87 | 0.87 | 0.86 | **0.880** | 0.870 | 0.007 |
| **SPAFESTWDILK** | SPAFESTWDILK | *Lead candidate* | (candidate) | 0.85 | 0.83 | 0.83 | 0.83 | 0.83 | **0.850** | 0.834 | 0.009 |
| **Binder group summary (n = 3)** | | | | **Mean best iPTM = 0.877 \| SD = 0.025 \| Range: 0.850 – 0.900** | | | | | **0.877** | 0.877 | 0.025 |
| **HARD NEGATIVES — 29 hard negatives across four categories (AF3 Server, 5 seeds, MDM2(25–109))** | | | | | | | | | | | |
| **ID / Peptide** | **Sequence** | **Category** | **Kd** | **Seed 1** | **Seed 2** | **Seed 3** | **Seed 4** | **Seed 5** | **Best iPTM** | **Mean iPTM** | **SD** |
| hn_w_pos_1 | WPAFESTDILK | *W-position* | - | 0.68 | 0.68 | 0.66 | 0.58 | 0.51 | **0.680** | 0.622 | 0.075 |
| hn_w_pos_2 | SWAFESTDILK | *W-position* | - | 0.43 | 0.36 | 0.37 | 0.35 | 0.32 | **0.430** | 0.366 | 0.040 |
| hn_w_pos_3 | SPWFESTDILK | *W-position* | - | 0.73 | 0.68 | 0.68 | 0.66 | 0.62 | **0.730** | 0.674 | 0.039 |
| hn_w_pos_4 | SPAFWSTDILK | *W-position* | - | 0.48 | 0.39 | 0.36 | 0.38 | 0.39 | **0.480** | 0.400 | 0.045 |
| hn_w_pos_5 | KLIDWTSEFAPS | *W-position* | - | 0.72 | 0.71 | 0.63 | 0.64 | 0.64 | **0.720** | 0.668 | 0.043 |
| hn_w_pos_6 | SPAFESTDIWK | *W-position* | - | 0.79 | 0.78 | 0.79 | 0.78 | 0.77 | **0.790** | 0.782 | 0.008 |
| hn_w_pos_7 | SPAFESTDILWK | *W-position* | - | 0.42 | 0.41 | 0.38 | 0.39 | 0.37 | **0.420** | 0.394 | 0.020 |
| hn_w_pos_8 | SPAFESTDIKLW | *W-position* | - | 0.35 | 0.32 | 0.33 | 0.37 | 0.29 | **0.370** | 0.332 | 0.029 |
| hn_scrm_wa_0 | TPASSLFIADEK | *Scrambled no-W* | - | 0.53 | 0.40 | 0.42 | 0.39 | 0.38 | **0.530** | 0.424 | 0.061 |
| hn_scrm_wa_1 | STASLIEDPAKF | *Scrambled no-W* | - | 0.46 | 0.47 | 0.42 | 0.38 | 0.37 | **0.470** | 0.420 | 0.044 |
| hn_scrm_wa_2 | PKILASEDSATF | *Scrambled no-W* | - | 0.39 | 0.36 | 0.35 | 0.37 | 0.35 | **0.390** | 0.364 | 0.016 |
| hn_scrm_wa_3 | DSLSFIPETAKA | *Scrambled no-W* | - | 0.36 | 0.38 | 0.36 | 0.35 | 0.35 | **0.380** | 0.360 | 0.012 |

| ID / Peptide | Sequence | Category | Kd | Seed 1 | Seed 2 | Seed 3 | Seed 4 | Seed 5 | Best iPTM | Mean iPTM | SD |
|---|---|---|---|---|---|---|---|---|---|---|---|
| hn_scrm_wa_4 | IPLATDSAESFK | *Scrambled no-W* | - | 0.41 | 0.34 | 0.42 | 0.33 | 0.30 | **0.420** | 0.360 | 0.051 |
| hn_hphob_nw_1 | SPAFESTDILK | *Hydrophobic no-W* | - | 0.35 | 0.33 | 0.31 | 0.31 | 0.34 | **0.350** | 0.328 | 0.018 |
| hn_hphob_nw_2 | LPAFESTDILK | *Hydrophobic no-W* | - | 0.43 | 0.37 | 0.35 | 0.37 | 0.37 | **0.430** | 0.378 | 0.031 |
| hn_hphob_nw_3 | SPAFESTVDILK | *Hydrophobic no-W* | - | 0.39 | 0.39 | 0.38 | 0.33 | 0.32 | **0.390** | 0.362 | 0.034 |
| hn_hphob_nw_4 | KPAFESTDILS | *Hydrophobic no-W* | - | 0.42 | 0.38 | 0.37 | 0.32 | 0.37 | **0.420** | 0.372 | 0.036 |
| hn_hphob_nw_5 | SPAFESTGILK | *Hydrophobic no-W* | - | 0.37 | 0.36 | 0.35 | 0.33 | 0.34 | **0.370** | 0.350 | 0.016 |
| hn_hphob_nw_6 | TPAFESDFILK | *Hydrophobic no-W* | - | 0.60 | 0.61 | 0.59 | 0.58 | 0.54 | **0.610** | 0.584 | 0.027 |
| hn_hphob_nw_7 | APSFETDILKV | *Hydrophobic no-W* | - | 0.43 | 0.41 | 0.45 | 0.34 | 0.38 | **0.450** | 0.402 | 0.042 |
| hn_hphob_nw_8 | FPASETDILKV | *Hydrophobic no-W* | - | 0.44 | 0.45 | 0.39 | 0.39 | 0.40 | **0.450** | 0.414 | 0.029 |
| hn_prot_rnd_1 | VGVNGFGRIGR | *Proteome-random* | - | 0.46 | 0.37 | 0.36 | 0.36 | 0.35 | **0.460** | 0.380 | 0.046 |
| hn_prot_rnd_2 | AGFAGDDAPR | *Proteome-random* | - | 0.40 | 0.39 | 0.35 | 0.34 | 0.39 | **0.400** | 0.374 | 0.026 |
| hn_prot_rnd_3 | INVYYNEATGGK | *Proteome-random* | - | 0.48 | 0.48 | 0.46 | 0.46 | 0.42 | **0.480** | 0.460 | 0.024 |
| hn_prot_rnd_4 | AILVDLEPGTMDS | *Proteome-random* | - | 0.55 | 0.46 | 0.44 | 0.37 | 0.37 | **0.550** | 0.438 | 0.075 |
| hn_prot_rnd_5 | DLYANTVLSGGTTM | *Proteome-random* | - | 0.38 | 0.36 | 0.40 | 0.32 | 0.34 | **0.400** | 0.360 | 0.031 |
| hn_prot_rnd_6 | VIHDHSGKVVK | *Proteome-random* | - | 0.29 | 0.23 | 0.22 | 0.22 | 0.20 | **0.290** | 0.232 | 0.035 |
| hn_prot_rnd_7 | EIAQDFKTDLR | *Proteome-random* | - | 0.47 | 0.46 | 0.47 | 0.45 | 0.43 | **0.470** | 0.456 | 0.016 |
| hn_prot_rnd_8 | LNVQNKDALLEK | *Proteome-random* | - | 0.41 | 0.39 | 0.39 | 0.36 | 0.33 | **0.410** | 0.376 | 0.031 |
| **Negative group summary (n = 29)** | | | | **Mean best iPTM = 0.474 \| SD = 0.122 \| Range: 0.290 – 0.790** | | | | | **0.474** | 0.474 | 0.122 |

*Matching criteria: W-position variants, scrambled no-W, and hydrophobic no-W categories were matched by amino acid composition to SPAFESTWDILK (same or analogous residues, lengths 11–12). Proteome-random peptides were filter-matched only (same six physicochemical filters as Supplementary Table S2; Boman index, GRAVY, charge, instability index, MW, hydrophobic fraction) and are not composition-matched; lengths range 10–14. Length was not strictly controlled across categories. Note: minimum gap between binder and negative best iPTM = +0.060 (SPAFESTWDILK 0.850 vs hn_w_pos_6 0.790); zero overlap across all 32 peptides. All per-seed values extracted from AF3 Server JSON files and cross-validated against Table S7.*

### Supplementary Table S4. EvoEF2 10-component inter-chain energy decomposition.

Full EvoEF2 inter-chain energy decomposition for the SPAFESTWDILK:MDM2(25–109) complex predicted by AlphaFold 3 Server (top-ranked model). Ten non-zero inter-chain (interD) energy terms are reported with physical descriptions, alongside all zero-value terms shown for completeness. Dominant contributions: inter-chain van der Waals attractive (−35.68 EEU [EvoEF2 Energy Units]), hydrophobic desolvation (−25.90 EEU), and polar desolvation penalty (+15.41 EEU). Total score: −47.27 EEU. The Protenix structure yields −55.57 EEU under identical settings. All intra-residue (intraR), intra-chain inter-residue (interS), and reference state terms are zero, confirming binding energy arises exclusively from inter-chain contacts.

*Technical note: the sum of the 10 non-zero interD terms equals −47.26 EEU while the JSON reports −47.27 EEU; the 0.01 discrepancy is a floating-point artefact from a signed-zero value (interD_hbscsc_phi = −0.0) in the EvoEF2 output. EvoEF2 job: neurosnap.ai/job/69df74bf. EEU units are not directly comparable to kcal/mol MM-PBSA values. Component breakdown for the Protenix structure (total −55.57 EEU): VdW attractive −39.16, hydrophobic desolvation −28.09 [h]; full decomposition available at Neurosnap job link [h].*

### Energy component decomposition (AF3 Server model_0, chain A vs chain B)

| EvoEF2 Term | Description | Physical meaning | Value (EEU) | Direction |
|---|---|---|---|---|
| **Non-zero inter-chain (interD) energy terms — interface contributions** | | | | |
| interD_vdwatt | Inter-chain VdW attractive | *Van der Waals attractive interactions between MDM2 and SPAFESTWDILK* | **-35.68** | *favourable* |
| interD_vdwrep | Inter-chain VdW repulsive | *Van der Waals steric clash / repulsion between chains* | **2.96** | *unfavourable* |
| interD_electr | Inter-chain electrostatics | *Electrostatic interactions between charged/polar groups across interface* | **-0.36** | *favourable* |
| interD_deslvP | Inter-chain desolvation (polar) | *Polar desolvation penalty upon complex formation* | **15.41** | *unfavourable* |
| interD_deslvH | Inter-chain desolvation (hydrophobic) | *Hydrophobic desolvation gain upon burial of hydrophobic surface* | **-25.90** | *favourable* |
| interD_hbscbb_dis | H-bond SC-BB distance | *Sidechain-to-backbone H-bond distance term* | **-1.61** | *favourable* |
| interD_hbscbb_the | H-bond SC-BB theta angle | *Sidechain-to-backbone H-bond angle term (theta)* | **-0.45** | *favourable* |
| interD_hbscbb_phi | H-bond SC-BB phi angle | *Sidechain-to-backbone H-bond angle term (phi)* | **-0.56** | *favourable* |
| interD_hbscsc_dis | H-bond SC-SC distance | *Sidechain-to-sidechain H-bond distance term* | **-0.92** | *favourable* |
| interD_hbscsc_the | H-bond SC-SC theta angle | *Sidechain-to-sidechain H-bond angle term (theta)* | **-0.15** | *favourable* |
| **TOTAL** | **Sum of all non-zero interD terms** | Overall predicted binding affinity (EvoEF2 score) | **-47.27** | *favourable* |
| **Zero-value terms (shown for completeness)** | | | | |
| *interD_hbscsc_phi* | H-bond SC-SC phi angle | *Not applicable / zero for this complex* | 0.00 | - |
| *interD_ssbond* | Inter-chain disulfide bond | *Not applicable / zero for this complex* | 0.00 | - |
| *interD_hbbbbb_dis* | H-bond BB-BB distance | *Not applicable / zero for this complex* | 0.00 | - |
| *interD_hbbbbb_the* | H-bond BB-BB theta | *Not applicable / zero for this complex* | 0.00 | - |
| *interD_hbbbbb_phi* | H-bond BB-BB phi | *Not applicable / zero for this complex* | 0.00 | - |

| EvoEF2 Term | Description | Physical meaning | Value (EEU) | Direction |
|---|---|---|---|---|
| *intraR_** | All intra-residue terms | *Not applicable / zero for this complex* | 0.00 | - |
| *interS_** | All intra-chain inter-residue terms | *Not applicable / zero for this complex* | 0.00 | - |
| *reference_** | Reference state energies (20 AA) | *Not applicable / zero for this complex* | 0.00 | - |

**Shading:** Green = favourable (stabilising); Amber = unfavourable (destabilising); White/Grey = zero (not applicable).

### EvoEF2 scores by structure input

| Structure input | Force field / method | EvoEF2 score (EEU) | Neurosnap job |
|---|---|---|---|
| AF3 Server model_0 | EvoEF2 (knowledge-based) | **-47.27** | 69df74bf |
| Protenix model_0 | EvoEF2 (knowledge-based) | **-55.57** | Neurosnap platform |

*Note: EvoEF2 scores depend on the input structure conformation. The Protenix structure (iPTM 0.923) scores more favourably than the AF3 structure (iPTM 0.850), consistent with its higher predicted confidence. Both structures predict strongly favourable binding (< -40 EEU). EEU units are not directly comparable to kcal/mol MM-PBSA values.*

*Technical note on arithmetic: the sum of the 10 non-zero interD terms listed above equals -47.26 EEU, whereas the EvoEF2 JSON reports TOTAL = -47.27 EEU. The 0.01 discrepancy arises because interD_hbscsc_phi is stored as -0.0 (a signed zero) in the output JSON, which contributes a sub-rounding floating-point value to the internal summation. This is a known floating-point artefact of the EvoEF2 implementation and does not represent an error in the table.*

### Supplementary Table S5. MM-PBSA binding free energy component breakdown.

Complete MM-PBSA binding free energy decomposition for SPAFESTWDILK:MDM2(25–109) under two independent force fields. Delta values represent binding contributions (ΔX = X_complex − X_receptor − X_ligand). AMBER99SB-ILDN applied to the Protenix structure: ΔG_total = −75.30 ± 4.92 kcal/mol (SD) across 12 reported energy terms (standard AMBER components). CHARMM27 applied to the AF3 structure: ΔG_total = −55.07 ± 2.86 kcal/mol (SD) across 15 reported energy terms; the three additional CHARMM27-specific bonded terms not present in AMBER are Urey–Bradley (ΔUB = 0.00 kcal/mol), improper dihedral (ΔIMP = +0.44 kcal/mol), and CMAP backbone correction (ΔCMAP = −1.42 kcal/mol). Electrostatics (ΔEEL) is the dominant favourable term in both force fields (−229.3 and −86.2 kcal/mol respectively); polar solvation (ΔEPB) is the dominant unfavourable counterterm. All values in kcal/mol; Avg = trajectory mean; SEM = standard error of mean across trajectory frames; SD = standard deviation across trajectory frames. Experimental ΔG reference values provided for PMI (Kd 3.3 nM, ΔG = −11.7 kcal/mol) and p53(17–29) (Kd ~580 nM, ΔG = −8.4 kcal/mol).

*MM-PBSA overestimates ΔG by 4–6× due to omission of the entropic term (−TΔS, estimated 10–30 kcal/mol for peptides of this size). Experimental ΔG values derived from Kd via ΔG = RT ln(Kd): PMI Kd = 3.3 nM (Pazgier et al. 2009); p53(17–29) Kd ~580 nM (Kussie et al. 1996). AMBER job: neurosnap.ai/job/69dfc05f. CHARMM27 job: neurosnap.ai/job/69e12743. Zenodo DOI: 10.5281/zenodo.19859570.*

#### Cross-force-field delta energy component comparison

| AMBER99SB-ILDN — Protenix structure \| ΔG_total = −75.30 ± 4.92 kcal/mol (SD) |
|---|

| Energy Component | Description | Avg (kcal/mol) | SEM | SD |
|---|---|---|---|---|
| ΔVDWAALS | *van der Waals* | -7.09 | 0.04 | 2.00 |
| ΔEEL | *Electrostatics* | -229.30 | 0.62 | 31.21 |
| Δ1-4 VDW | *1-4 van der Waals* | 0.90 | 0.01 | 0.44 |
| Δ1-4 EEL | *1-4 Electrostatics* | 61.75 | 0.05 | 2.27 |
| ΔBOND | *Bond* | 0.31 | 0.01 | 0.44 |
| ΔANGLE | *Angle* | 1.06 | 0.02 | 0.83 |
| ΔDIHED | *Dihedral* | 7.75 | 0.02 | 1.14 |
| ΔEPB | *Polar solvation (EPB)* | 162.34 | 0.55 | 27.41 |
| ΔENPOLAR | *Non-polar solvation* | -1.25 | 0.00 | 0.19 |
| ΔGGAS | *Gas-phase total (GGAS)* | -236.39 | 0.62 | 30.97 |
| ΔGSOLV | *Solvation total (GSOLV)* | 161.09 | 0.55 | 27.34 |
| **ΔTOTAL** | TOTAL DeltaG | **-75.30** | 0.10 | 4.92 |

**CHARMM27 — AF3 structure | ΔG_total = −55.07 ± 2.86 kcal/mol (SD) | includes 3 CHARMM27-specific terms (amber shading)**

| Energy Component | Description | Avg (kcal/mol) | SEM | SD |
|---|---|---|---|---|
| ΔVDWAALS | *van der Waals* | -2.36 | 0.03 | 1.46 |
| ΔEEL | *Electrostatics* | -86.21 | 0.51 | 25.75 |
| Δ1-4 VDW | *1-4 van der Waals* | 0.86 | 0.01 | 0.51 |
| Δ1-4 EEL | *1-4 Electrostatics* | 35.36 | 0.03 | 1.70 |
| ΔBOND | *Bond* | 0.32 | 0.01 | 0.45 |
| ΔANGLE | *Angle* | 1.12 | 0.02 | 0.90 |
| ΔDIHED | *Dihedral* | 2.39 | 0.01 | 0.67 |
| ΔUB | *Urey-Bradley (CHARMM27 only)* | 0.00 | 0.00 | 0.00 |
| ΔIMP | *Improper dihedral (CHARMM27 only)* | 0.44 | 0.01 | 0.47 |
| ΔCMAP | *CMAP correction (CHARMM27 only)* | -1.42 | 0.02 | 0.85 |
| ΔEPB | *Polar solvation (EPB)* | 35.30 | 0.47 | 23.47 |
| ΔENPOLAR | *Non-polar solvation* | -0.83 | 0.00 | 0.10 |
| ΔGGAS | *Gas-phase total (GGAS)* | -89.54 | 0.50 | 25.23 |
| ΔGSOLV | *Solvation total (GSOLV)* | 34.47 | 0.47 | 23.38 |
| **ΔTOTAL** | TOTAL DeltaG | **-55.07** | 0.06 | 2.86 |

*Note on CHARMM27-specific terms: CHARMM27 includes three additional bonded energy terms not present in AMBER99SB-ILDN — Urey-Bradley (UB, harmonic angle potential acting on 1-3 atom pairs), improper dihedral (IMP, planarity constraint), and CMAP (backbone phi/psi cross-map correction). UB = 0.00, IMP = +0.44, CMAP = -1.42 kcal/mol for this complex; their combined contribution is -0.98 kcal/mol. These terms partially account for the force-field difference in TOTAL DeltaG (-55.07 vs -75.30 kcal/mol), with the dominant contribution coming from the EEL term (-86.21 vs -229.30 kcal/mol).*

**Shading key:** Green = favourable (negative delta, stabilising binding); Amber = unfavourable (positive delta, destabilising); Grey = near zero or sub-term.

### Experimental DeltaG reference values (for context)

| Reference compound | Kd | Experimental ΔG (kcal/mol) | Source |
|---|---|---|---|
| PMI | 3.3 nM | -11.7 | RT ln(Kd); Pazgier et al. 2009 |
| p53(17-29) | ~580 nM | -8.4 | RT ln(Kd); Kussie et al. 1996 |
| SPAFESTWDILK (predicted) | - | - | No experimental Kd available |

*Note: Experimental ΔG derived from Kd via ΔG = RT ln(Kd). PMI: Kd = 3.3 nM (Pazgier et al. 2009). p53(17–29): Kd ~580 nM (Kussie et al. 1996). TOTAL MM-PBSA values are 4–6× larger in magnitude than experimental ΔG, consistent with known systematic overestimation by implicit-solvent MM-PBSA in the absence of −TΔS correction.*

### Supplementary Table S6. ExPASy PeptideCutter: GI proteolytic digestion analysis.

In silico proteolytic digestion of SPAFESTWDILK using 13 GI-relevant enzyme specificities from ExPASy PeptideCutter (Gasteiger et al. 2005). Enzymes cover the major GI proteolytic environment: gastric (pepsin pH 1.3 and >2), pancreatic (trypsin, chymotrypsin high/low specificity, elastase, pancreatic elastase II), intestinal (Glu-C/V8 pH 8.0 and 4.0, proline endopeptidase), and plant/microbial (bromelain, lysyl endopeptidase, clostripain, thrombin). For each enzyme: cleavage rule, number and positions of cleavage sites, fragments produced, GI phase, and susceptibility classification. Cleavage rules follow official ExPASy documentation: elastase cleaves after A, G, S, V only (T is not a substrate); proline endopeptidase cleaves after P (P-X bonds); Glu-C/V8 is pH-dependent (pH 8.0: after E only; pH 4.0: after E and D). The two primary MDM2 pharmacophore anchors F4 and W8 are cleavage sites for pepsin and chymotrypsin, indicating susceptibility to gastric and pancreatic proteolysis. Terminal K12 cleavage by trypsin, Lys-C, and bromelain produces the intact peptide as the only fragment and is classified as functionally resistant. A residue-level annotation table maps each position to enzymes, GI phase, and MDM2-binding role.

*All rules verified against ExPASy PeptideCutter documentation (expasy.org/resources/peptidecutter). GI susceptibility at F4 and W8 is a standard limitation for linear dietary peptides and does not preclude MDM2 activity at sites of local absorption or under protective formulation.*

#### Enzyme cleavage summary

| Enzyme | Cleavage rule | N sites | Site positions | Fragments produced | GI context | Susceptibility |
|---|---|---|---|---|---|---|
| **GI-relevant proteolytic enzymes** | | | | | | |
| **Pepsin (pH 1.3)** | *Cleaves after F, L, W; not before P* | **3** | F4, W8, L11 | SPAF \| ESTW \| DIL \| K | *Gastric (pH 1.3–2.0)* | No — 3 sites |
| **Pepsin (pH > 2)** | *Cleaves after F, L; not before P* | **2** | F4, L11 | SPAF \| ESTWDIL \| K | *Gastric (pH > 2.0)* | No — 2 sites |
| **Trypsin** | *Cleaves after K, R; not before P* | **1** | K12 (terminal) | **SPAFESTWDILK (intact)** | *Pancreatic / intestinal* | **Functionally resistant*** |
| **Chymotrypsin (high specificity)** | *Cleaves after F, Y, W; not before P* | **2** | F4, W8 | SPAF \| ESTW \| DILK | *Pancreatic* | No — 2 sites |

| Enzyme | Cleavage rule | N sites | Site positions | Fragments produced | GI context | Susceptibility |
|---|---|---|---|---|---|---|
| **Chymotrypsin (low specificity)** | *Cleaves after F, Y, W, L, M; not before P* | **3** | F4, W8, L11 | SPAF \| ESTW \| DIL \| K | *Pancreatic* | No — 3 sites |
| **Elastase** | *Cleaves after A, G, S, V; not before P (T is NOT a substrate)* | **2** | A3, S6 | SPA \| FES \| TWDILK | *Pancreatic* | No — 2 sites |
| **Pancreatic elastase II** | *Cleaves after L, I, V; not before P* | **2** | I10, L11 | SPAFESTWDI \| L \| K | *Pancreatic* | No — 2 sites |
| **Bromelain** | *Cleaves after K, R; not before P* | **1** | K12 (terminal) | **SPAFESTWDILK (intact)** | *Plant cysteine protease (ginger source)* | **Functionally resistant*** |
| **Proline endopeptidase** | *Cleaves AFTER P (P-X bonds where X is not P)* | **1** | P2 (cleaves SP\|AFESTWDILK) | SP \| AFESTWDILK | *Intestinal (post-Pro sequences)* | No — 1 site |
| **Glutamyl endopeptidase (V8/GluC) pH 8.0** | *Cleaves after E only at pH 8.0* | **1** | E5 | SPAFE \| STWDILK | *Intestinal (pH 8, minor)* | No — 1 site |
| **Glutamyl endopeptidase (V8/GluC) pH 4.0** | *Cleaves after E and D at pH 4.0* | **2** | E5, D9 | SPAFE \| STWD \| ILK | *Intestinal (pH 4, minor)* | No — 2 sites |
| **Lysyl endopeptidase (Lys-C)** | *Cleaves after K* | **1** | K12 (terminal) | **SPAFESTWDILK (intact)** | *Lysosomes / gut microbiome* | **Functionally resistant*** |
| **Thrombin** | *Cleaves R-G bonds* | **0** | None | **SPAFESTWDILK (intact)** | *Coagulation (not GI)* | Yes — no sites |
| **Clostripain** | *Cleaves after R* | **0** | None | **SPAFESTWDILK (intact)** | *Anaerobic gut bacteria* | Yes — no sites |

### Residue-level annotation — cleavage sites mapped to MDM2-binding pharmacophore

| Position | Residue | Enzymes cleaving here | GI phase | MDM2-binding role |
|---|---|---|---|---|
| **1** | **S (Ser)** | *No enzyme — S1 not cleaved* | — | N-terminal; not in MDM2 contact |
| **2** | **P (Pro)** | *Proline endopeptidase (cleaves AFTER P2 -> SP\|AFESTWDILK)* | Intestinal | Not in MDM2 contact |
| **3** | **A (Ala)** | *Elastase (A,G,S,V rule — no T)* | Pancreatic | Not in MDM2 contact |
| **4** | **F (Phe)** | *Pepsin (pH 1.3 and >2); Chymotrypsin (high+low spec.)* | Gastric + pancreatic | **Phe19-equivalent — MDM2 Phe19 sub-pocket anchor** |
| **5** | **E (Glu)** | *Glu-C/V8 (pH 8.0 and 4.0)* | Intestinal (minor) | Not in MDM2 direct contact |
| **6** | **S (Ser)** | *Elastase (A,G,S,V rule)* | Pancreatic | Not in MDM2 contact |
| **7** | **T (Thr)** | *No enzyme (T not in elastase or other GI rule)* | — | Not in MDM2 contact |

| Position | Residue | Enzymes cleaving here | GI phase | MDM2-binding role |
|---|---|---|---|---|
| **8** | **W (Trp)** | *Pepsin (pH 1.3); Chymotrypsin (high+low spec.)* | Gastric + pancreatic | **Trp23-equivalent — principal MDM2 anchor** |
| **9** | **D (Asp)** | *Glu-C/V8 (pH 4.0 only)* | Intestinal (minor, pH 4.0 only) | Not in MDM2 direct contact |
| **10** | **I (Ile)** | *Pancreatic elastase II* | Pancreatic | Leu26 sub-pocket (joint with L11) |
| **11** | **L (Leu)** | *Pepsin (pH 1.3); Chymotrypsin (low spec.); Pancreatic elastase II* | Gastric + pancreatic | **Leu26 sub-pocket anchor** |
| **12** | **K (Lys)** | *Trypsin; Lys-C; Bromelain (terminal — intact peptide produced)* | Tryptic release site | C-terminal; confirms tryptic origin |

*Key finding: F4 and W8 — the two primary MDM2 pharmacophore anchors — are cleavage sites for pepsin (gastric) and chymotrypsin (pancreatic). This means SPAFESTWDILK is susceptible to proteolysis at its functionally critical residues under GI conditions. L11 (Leu26 sub-pocket) is also a chymotrypsin/elastase site. The peptide retains its full sequence only if absorbed intact (e.g. via paracellular transport or in a formulation that protects it from luminal proteases). This is a standard limitation for linear food-derived peptides and is explicitly acknowledged in the manuscript.*

### Supplementary Table S7. Effect size derivation: Cohen's d and Hedges' g.

Full derivation of Cohen's d, Hedges' g, and 95% confidence intervals for the 36-peptide AF3 benchmark (Panel A) and the seven-peptide Protenix discrimination panel (Panel B). Panel A: binder group (n = 3: PMI, 12/1, SPAFESTWDILK; mean best iPTM = 0.877, SD = 0.025) vs hard-negative group (n = 29; mean best iPTM = 0.474, SD = 0.122). Effect sizes computed on individual peptide best-iPTM values using df-weighted pooled SD: pooled SD = $\sqrt{[((n_1-1)s_1^2 + (n_2-1)s_2^2) / (n_1+n_2-2)]}$. Cohen's d = 3.41 (95% CI: 1.94–4.88); Hedges' g = 3.32 (correction factor J = 0.9748). Zero overlap confirmed: minimum binder best-iPTM (0.850) exceeds maximum negative best-iPTM (0.790) by +0.060. Panel B: binder group (n = 3; mean iPTM = 0.928, SD = 0.011) vs non-binder group (n = 4; mean iPTM = 0.741, SD = 0.037). Cohen's d = 6.43; Hedges' g = 5.42 (descriptive only, small n). W8A/W23A mirror: Δ(SPAFESTWDILK→W8A) = 0.201; Δ(p53→W23A) = 0.193; difference = 0.008.

*95% CI for Cohen's d uses SE(d) = $\sqrt{[N/(n_1 n_2) + d^2/(2N)]}$. All iPTM values in Panel A are AF3 Server best-seed scores; Panel B values are Protenix best-seed scores. Raw data in AF3_Server_Benchmark_29_Hard_Negatives.zip and AF3_Server_Controls_8peptides_5seeds.zip (Zenodo DOI: 10.5281/zenodo.19859570).*

#### Panel A: 36-Peptide Expanded AF3 Benchmark

Known binders (n = 3: PMI, 12/1, SPAFESTWDILK) vs 29 hard negatives across four categories. All evaluations on AF3 Server, 5 seeds per peptide, MDM2(25–109) construct.

| ID / Peptide | Category / Kd | Sequence / Seeds (5 values) | Best iPTM | Avg iPTM | SD |
|---|---|---|---|---|---|
| **BINDERS — AF3 Server best-seed iPTM (5 seeds each)** | | | | | |
| **PMI** | 3.3 nM | 0.90, 0.89, 0.89, 0.89, 0.88 | **0.900** | 0.890 | 0.0071 |
| **12/1** | ~200 nM | 0.88, 0.87, 0.87, 0.87, 0.86 | **0.880** | 0.870 | 0.0071 |
| **SPAFESTWDILK** | (candidate) | 0.85, 0.83, 0.83, 0.83, 0.83 | **0.850** | 0.834 | 0.0089 |
| **Binder group summary (n = 3)** | | **Mean best iPTM = 0.877** | SD = 0.025 | N = 3 | |
| **HARD NEGATIVES — AF3 Server best-seed iPTM (29 peptides)** | | | | | |
| **ID** | **Category** | **Sequence** | **Best iPTM** | | |

| ID / Peptide | Category / Kd | Sequence / Seeds (5 values) | Best iPTM | Avg iPTM | SD |
|---|---|---|---|---|---|
| hn_w_pos_1 | *W-position* | WPAFESTDILK | **0.680** | | |
| hn_w_pos_2 | *W-position* | SWAFESTDILK | **0.430** | | |
| hn_w_pos_3 | *W-position* | SPWFESTDILK | **0.730** | | |
| hn_w_pos_4 | *W-position* | SPAFWSTDILK | **0.480** | | |
| hn_w_pos_5 | *W-position* | KLIDWTSEFAPS | **0.720** | | |
| hn_w_pos_6 | *W-position* | SPAFESTDIWK | **0.790** | | |
| hn_w_pos_7 | *W-position* | SPAFESTDILWK | **0.420** | | |
| hn_w_pos_8 | *W-position* | SPAFESTDIKLW | **0.370** | | |
| hn_scrm_wa_0 | *Scrambled no-W* | TPASSLFIADEK | **0.530** | | |
| hn_scrm_wa_1 | *Scrambled no-W* | STASLIEDPAKF | **0.470** | | |
| hn_scrm_wa_2 | *Scrambled no-W* | PKILASEDSATF | **0.390** | | |
| hn_scrm_wa_3 | *Scrambled no-W* | DSLSFIPETAKA | **0.380** | | |
| hn_scrm_wa_4 | *Scrambled no-W* | IPLATDSAESFK | **0.420** | | |
| hn_hphob_nw_1 | *Hydrophobic no-W* | SPAFESTDILK | **0.350** | | |
| hn_hphob_nw_2 | *Hydrophobic no-W* | LPAFESTDILK | **0.430** | | |
| hn_hphob_nw_3 | *Hydrophobic no-W* | SPAFESTVDILK | **0.390** | | |
| hn_hphob_nw_4 | *Hydrophobic no-W* | KPAFESTDILS | **0.420** | | |
| hn_hphob_nw_5 | *Hydrophobic no-W* | SPAFESTGILK | **0.370** | | |
| hn_hphob_nw_6 | *Hydrophobic no-W* | TPAFESDFILK | **0.610** | | |
| hn_hphob_nw_7 | *Hydrophobic no-W* | APSFETDILKV | **0.450** | | |
| hn_hphob_nw_8 | *Hydrophobic no-W* | FPASETDILKV | **0.450** | | |
| hn_prot_rnd_1 | *Proteome-random* | VGVNGFGRIGR | **0.460** | | |
| hn_prot_rnd_2 | *Proteome-random* | AGFAGDDAPR | **0.400** | | |
| hn_prot_rnd_3 | *Proteome-random* | INVYYNEATGGK | **0.480** | | |
| hn_prot_rnd_4 | *Proteome-random* | AILVDLEPGTMDS | **0.550** | | |
| hn_prot_rnd_5 | *Proteome-random* | DLYANTVLSGGTTM | **0.400** | | |
| hn_prot_rnd_6 | *Proteome-random* | VIHDHSGKVVK | **0.290** | | |
| hn_prot_rnd_7 | *Proteome-random* | EIAQDFKTDLR | **0.470** | | |
| hn_prot_rnd_8 | *Proteome-random* | LNVQNKDALLEK | **0.410** | | |
| **Negative group summary (n = 29)** | | **Mean best iPTM = 0.474** | SD = 0.122 | N = 29 | |

### Panel A — Effect Size Calculation

| Parameter | Value / Formula |
|---|---|
| **Formula** | $d = (\bar{x}\_binders - \bar{x}\_neg) / pooled\_SD$ (df-weighted) |
| **Mean binders ($\bar{x}_1$)** | 0.877 |
| **Mean negatives ($\bar{x}_2$)** | 0.474 |
| **Var binders ($s_1^2$)** | 0.000634 (SD = 0.0252) |
| **Var negatives ($s_2^2$)** | 0.014953 (SD = 0.1223) |
| **Pooled SD** | $\sqrt{[((n_1-1)\cdot s_1^2 + (n_2-1)\cdot s_2^2) / (n_1+n_2-2)]} = \sqrt{[(2\times0.000634 + 28\times0.014953) / 30]} = 0.1183$ |
| **Cohen's d** | (0.877 − 0.474) / 0.1183 = 3.41 |
| **Correction factor J** | 1 − 3 / (4·df − 1) = 1 − 3/119 = 0.9748 |
| **Hedges' g** | 3.41 × 0.9748 = 3.32 |
| **SE(d)** | $\sqrt{[N/(n_1\cdot n_2) + d^2/(2\cdot N)]} = \sqrt{[32/87 + 11.63/64]} = 0.7411$ |
| **95% CI** | 3.41 ± 1.96 × 0.7411 = [1.94, 4.88] |
| **Zero overlap** | min(binders) = 0.850 > max(negatives) = 0.790 → gap = +0.060 |

*Note: Cohen's d computed on individual best-iPTM values (3 binders + 29 negatives = 32 total), not on group means. Pooled SD is df-weighted: $[(n_1-1)s_1^2 + (n_2-1)s_2^2] / (n_1+n_2-2)$. Hedges' g correction factor $J = 1 - 3/(4df-1)$ where $df = n_1+n_2-2 = 30$. $SE(d) = \sqrt{[N/(n_1\cdot n_2) + d^2/(2N)]}$ where $N = n_1+n_2$. Zero overlap confirmed: all 3 binder best-iPTM values (0.850–0.900) exceed all 29 negative best-iPTM values (0.290–0.790).*

### Panel B: 7-Peptide Protenix Discrimination Panel

Binders (n = 3) vs non-binders (n = 4). All evaluations on Neurosnap Protenix platform, MDM2(25–109) construct, 5 models each. W8A and W23A mirror effect is the study's principal falsifiable prediction.

| Peptide | Protenix iPTM | Class | Neurosnap Job ID |
|---|---|---|---|
| **PMI (3.3 nM)** | **0.940** | *Binder* | 69df89d5 |
| **SPAFESTWDILK** | **0.923** | *Binder* | 69df686c |
| **p53(15-29) (~140 nM)** | **0.920** | *Binder* | 69df89bb |
| Poly-Ala (ctrl−) | **0.795** | *Non-binder* | 69df8844 |
| p53 W23A (ctrl−) | **0.727** | *Non-binder* | 69df89f1 |
| SPAFESTADILK W8A (ctrl−) | **0.722** | *Non-binder* | 69df8956 |
| Scrambled (ctrl−) | **0.718** | *Non-binder* | 69df88ac |
| **Binder group: mean = 0.928, SD = 0.011, n = 3** | | **Non-binder group: mean = 0.741, SD = 0.036, n = 4** | |

### Panel B — Effect Size Calculation

| Parameter | Value / Formula |
|---|---|
| **Mean binders ($\bar{x}_1$)** | 0.9277 |
| **Mean non-binders ($\bar{x}_2$)** | 0.7405 |
| **SD binders ($s_1$)** | 0.0108 |
| **SD non-binders ($s_2$)** | 0.0365 |
| **Pooled SD** | $\sqrt{[((3-1)\cdot0.0108^2 + (4-1)\cdot0.0365^2) / (3+4-2)]} = 0.0291$ |
| **Cohen's d** | (0.9277 − 0.7405) / 0.0291 = 6.43 |

| Parameter | Value / Formula |
|---|---|
| **Hedges' g** | 6.43 × (1 − 3/(4·5−1)) = 5.42 |
| **95% CI (d)** | [2.75, 10.12] — wide due to n=3 vs n=4 |
| **W8A drop (SPAFESTWDILK)** | 0.923 − 0.722 = 0.201 |
| **W23A drop (p53)** | 0.920 − 0.727 = 0.193 |
| **Mirror difference** | \|0.201 − 0.193\| = 0.008 → approximately equal |

*Note: With n = 3 vs n = 4, Cohen's d = 6.43 and Hedges' g = 5.42 are descriptive only — the 95% CI spans [2.75, 10.12] reflecting the small n. The W8A/W23A mirror (Δ = 0.201 vs 0.193, difference = 0.008) is the study's principal testable hypothesis: Trp8 of SPAFESTWDILK is predicted to occupy the MDM2 Trp23 sub-pocket analogously to Trp23 of p53.*
